# An Integrated Atlas of the Human Kidney Spanning Health and Disease

**DOI:** 10.64898/2026.08.12.744548

**Authors:** Konstantinos Stasinos, Hanchen Wang, Alexander V. Predeus, Nathan Richoz, Rajasree Menon, Mrunali Thokadiwala, Yuefei Zhu, Ruijia Tian, Wenjiang Zhou, Arsenios Chatzigeorgiou, Galabina Yordanova, Ida Zucchi, Zoltan Laszik, Michaela F. Mueller, The Human Cell Atlas Kidney Bionetwork, Ayshwarya Subramanian, Anna Greka, Aviv Regev, Matthias Kretzler, John Marioni, Malte D. Luecken, Menna Clatworthy, Sarah A. Teichmann, Peng He

**Affiliations:** Department of Pathology, ImmunoX, University of California, San Francisco, CA, USA; Department of Surgery, University of Cambridge, UK; Department of Engineering, University of Cambridge, UK; Genentech, South San Francisco, CA, USA; Wellcome Sanger Institute, Wellcome Genome Campus, UK; Molecular Immunity Unit, MRC Laboratory of Molecular Biology, University of Cambridge, UK; Michigan Medicine (University of Michigan), Department of Computational Medicine & Bioinformatics, Ann Arbor, MI, USA; Department of Molecular Biology and Genetics, College of Arts & Sciences, Cornell University, Ithaca, NY, USA; European Molecular Biology Laboratory, European Bioinformatics Institute (EMBL-EBI), Wellcome Genome Campus Hinxton UK; Institute of Computational Biology, Helmholtz Munich, Neuherberg, Germany; Harvard Medical School, Boston, MA, USA; Brigham and Women’s Hospital, Boston, MA, USA; Cancer Research UK Cambridge Institute, University of Cambridge, UK; Institute of Lung Health and Immunity (LHI), Helmholtz Munich, Comprehensive Pneumology Center (CPC-M), Germany; Member of the German Center for Lung Research (DZL); Department of Medicine, University of Cambridge, UK; Cambridge Stem Cell Institute, University of Cambridge, UK; CIFAR Macmillan Multi-scale Human Programme, CIFAR, Toronto, Canada

## Abstract

The human kidney contains highly specialized cell populations. Despite numerous single-cell and single-nucleus transcriptomics studies, differences in cohorts, technologies, analytical pipelines, and annotation frameworks have limited the ability to define consensus kidney cell states, identify disease-associated populations and interpret kidney disease genetic susceptibility. Here, we assembled 18 human kidney single-cell and single-nucleus RNA-sequencing datasets spanning 232 donors and five major disease contexts into a uniformly processed and computationally integrated Human Kidney Cell Atlas (HKCA), comprising over one million high-quality cells (816,895) and nuclei (215,308). The HKCA resolves 63 cell types and 120 harmonized cell states, including rare epithelial and stromal populations associated with kidney disease. Integration with spatial transcriptomics, intercellular communication networks, and human genetic association data further defined the anatomical context and disease relevance of these populations. The HKCA also provides a framework for automated annotation of independent human and mouse kidney datasets. Together, the HKCA establishes a comprehensive reference for human kidney biology, enabling disease interpretation and genetic risk localization at cellular resolution.

## Main

The kidney maintains systemic homeostasis through a highly organized architecture of glomerular, tubular, vascular, stromal, immune and neural-associated compartments. Each nephron segment carries out specialized transport and endocrine functions, while stromal, endothelial and immune populations shape tissue architecture, oxygen sensing, inflammatory responses and repair.^1,2^ Inherited or acquired disruption of specific kidney cell populations contributes to clinically diverse diseases, including acute kidney injury^3–5^, diabetic kidney disease^6–8^, immune-mediated nephritis^9,10^, polycystic kidney disease^11^ and transplant rejection^12,13^ which can lead to long-term functional decline described as chronic kidney disease (CKD) which is estimated to affect 788 million people worldwide.^14–16^ Similarly, genetic susceptibility is increasingly recognized as an important contributor to both inherited and common kidney diseases; however, our understanding of how genetic variation influences kidney biology, particularly at the level of specific renal cell types and molecular pathways, remains incomplete.^7^

Single-cell and single-nucleus RNA-sequencing (sc/snRNA-seq) studies have transformed our understanding of kidney biology by resolving major nephron segments, immune populations, endothelial subtypes and stromal compartments in human tissue. However, existing datasets differ in donor composition, disease representation, sample handling, dissociation protocol, sequencing modality, genome alignment, ambient RNA correction, clustering strategy and annotation vocabulary. As a result, kidney cell-state nomenclature remains inconsistent across studies (Ext. Data Fig. 1), and subtle or rare populations are often difficult to distinguish from study-specific or modality-specific artifacts.^17,18^

**Figure 1:**
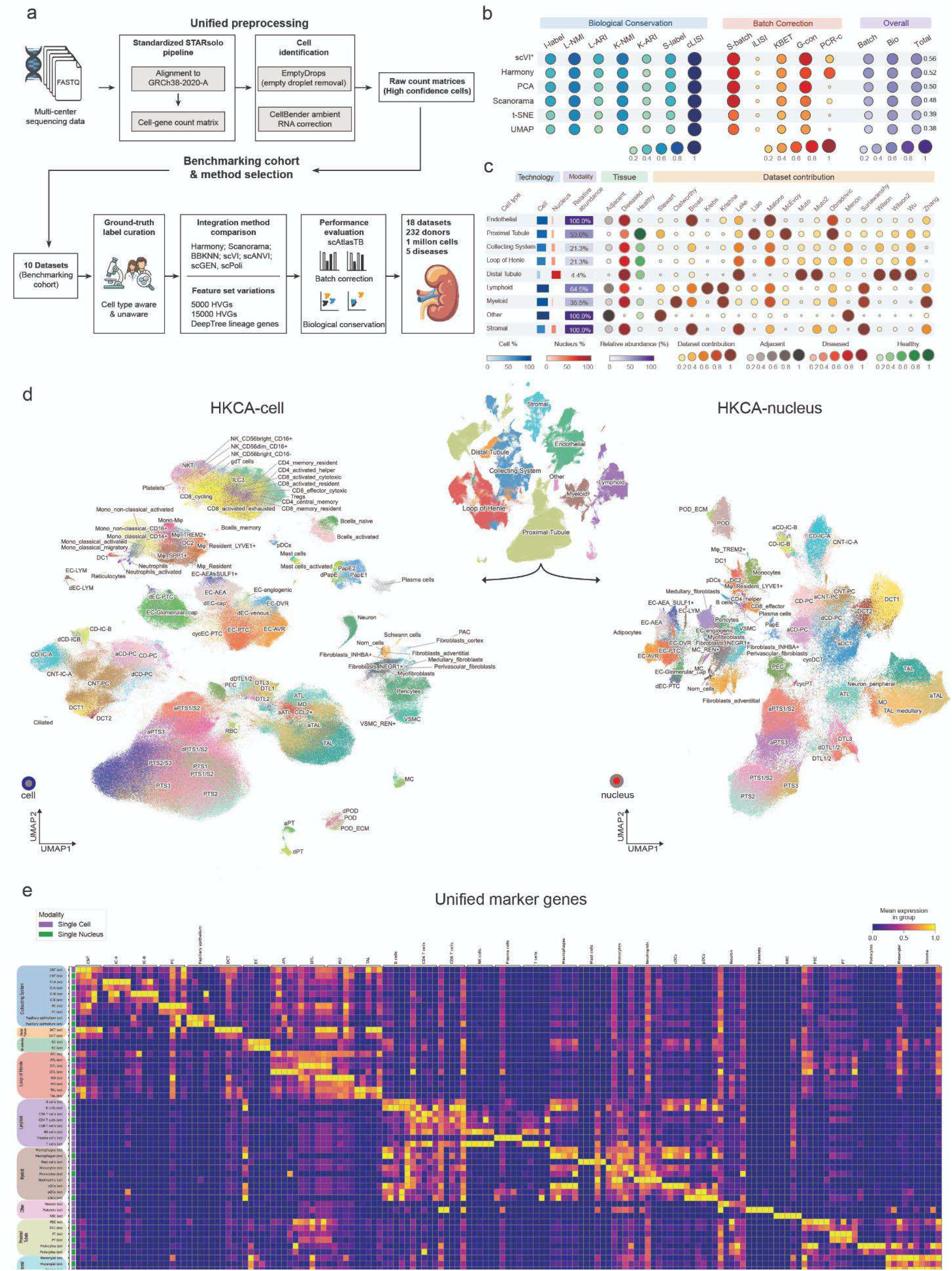
Constructing the Human Kidney Cell Atlas. a)18 individual datasets were reprocessed by accumulating the individual FASTQ files and remapped to a common reference genome using STARsolo (see Methods for detailed preprocessing). A subset of these datasets used as a core dataset was manually annotated and harmonized with CellHint and manual curation to provide a supervised benchmarking for subsequent integration metrics. The final objects compromise 1,032,203 cells and nuclei from 232 donors, spanning normal and disease states. b) Integration metrics for the ∼15,786 Deeptree^43^ algorithm selected genes across single cell and nucleus with scVI outperforming in the summative integration metrics. (See Ext. Fig. 2 for results of other integration settings) c) Composition of the atlas across cell types based on single cell and nucleus RNA sequencing and dataset contribution. d, e) The HKCA-cell and HKCA-nucleus integrated objects comprising the HKCA after isolating the modality specific datasets and the unified marker selection heatmap across both modalities based on the union of identified markers and scaled per modality.

A consensus human kidney cell atlas must therefore solve two related problems. First, it must standardize computational processing across studies so that differences in preprocessing such as intron inclusion or not, reference transcriptome version and alignment methods, do not masquerade as biological variation. Second, it must incorporate single-cell and single-nucleus data which have profound modality-specific different dissociation sensitivity, nuclear transcript content and susceptibility to ambient RNA contamination.^19^ Initial efforts to standardize kidney sample collection start to provide important insights into kidney biology and disease states.^16,20^ Nevertheless, they have been concentrated on a single modality, and geographically confined populations which might have moderate genomic variability.

To address these challenges, we assembled a comprehensive, modality-aware Human Kidney Cell Atlas (HKCA) by integrating human kidney single-cell and single-nucleus RNA-seq datasets across diverse studies, donors, and disease contexts using a unified computational framework.

## Results

### Construction of a uniformly processed HKCA across health and disease

We first assembled a compendium of human kidney sn/scRNA-seq datasets selected to maximize donor diversity, disease representation and availability of raw sequencing data. The final collection included **18 datasets,** ^6,7,9,11,12,16,21–30^ **232 donors**, healthy and adjacent-normal kidney tissues (n=195), and five major disease contexts: **acute kidney injury (**n=16)^16^**, immune-mediated nephritis (**n=2)^9^**, transplant-associated disease (**n=6)^12^**, diabetic kidney disease (**n=5)^7^ **and autosomal dominant polycystic kidney disease (**n=8)^11^ (Ext. Data Fig. 2c, 3, 4).

**Figure 2.**
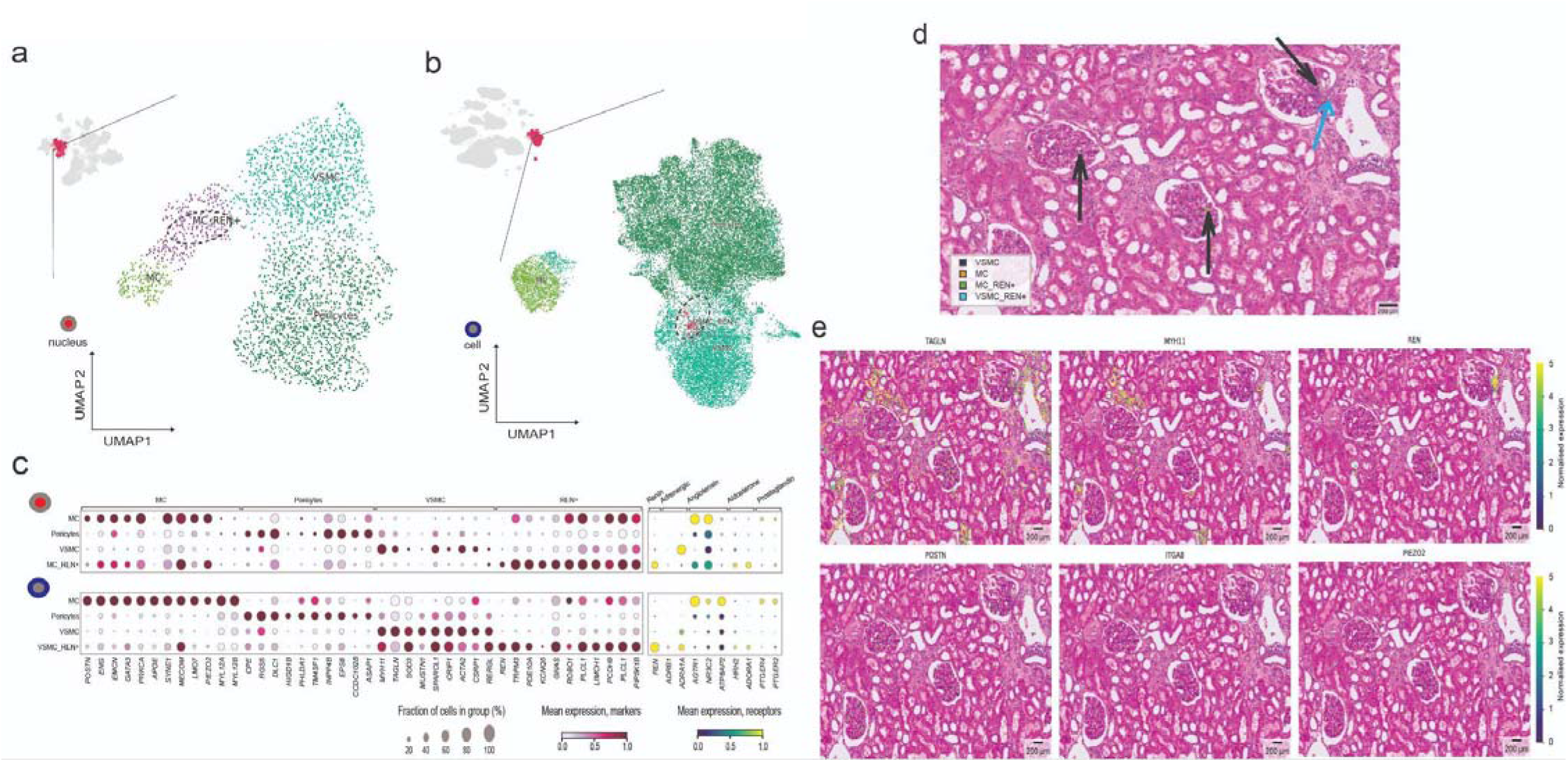
Renin is produced by discrete populations of mural cells. a,b) UMAP embeddings of stromal populations from the HKCA-nucleus (part a) or HKCA-cell (part b) highlighting renin-expressing subsets within vascular smooth muscle cells (VSMC-REN□) and mesangial cells (MC-REN□), indicating transcriptionally distinct renin-producing populations. c) Dot plot of marker gene expression across VSMC and mesangial populations. *REN* is expressed in both VSMC-REN and MC-REN subsets, alongside lineage markers (e.g., *MYH11*, *ACTA2* for VSMCs; *PIEZO2* for mesangial cells) for the HKCA-nucleus (top) and HKCA-cell (bottom). Also shown is the expression pattern of renin controlling receptors. d) Spatial transcriptomics (Visium HD) showing localization of REN-expressing cells to both vascular and glomerular compartments (arrows). e) Spatial expression of VSMC markers (TAGLN, MYH11), matrix-associated genes (POSTN, ITGA8), and REN.

**Figure 3.**
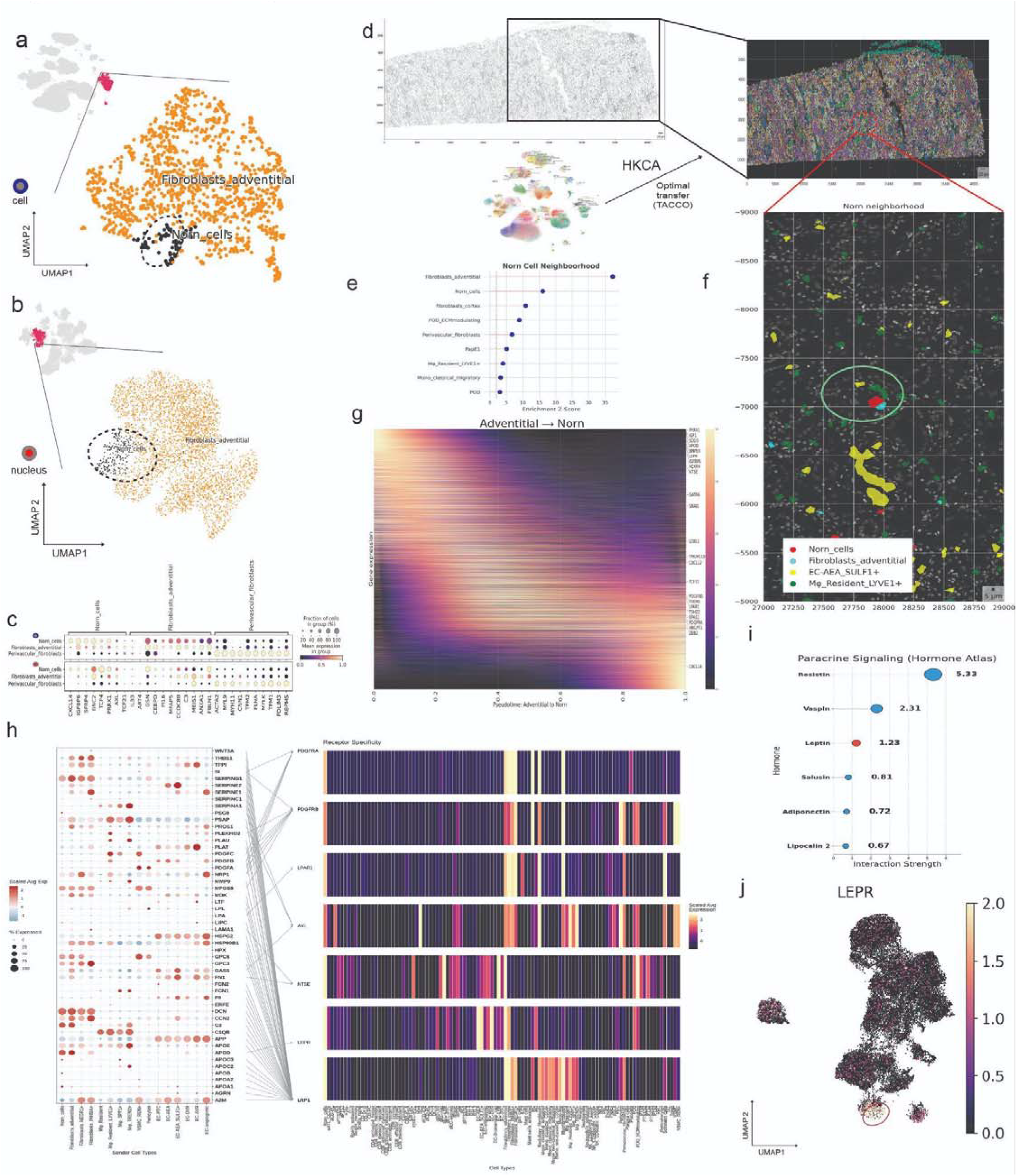
Norn cells reside in perivascular niches. a,b) UMAP of stromal cells from the HKCA highlighting Norn cells across HKCA-cell(part a) and HKCA-nucleus (part b) datasets (dashed circle). c) Dot plots comparing adventitial and perivascular fibroblast gene expression with Norn cell gene expression in the HKCA-cell (top) and HKCA-nucleus (bottom). d) Schematic of HKCA-guided annotation of Xenium spatial transcriptomics data using optimal transport (TACCO). Cell type identities from the HKCA are mapped onto spatial gene expression profiles and visualized across the tissue section. e) Neighborhood enrichment analysis [Enrichment Z-scores (squidpy)] showing spatial associations of Norn cells with surrounding cell types. f) Spatial visualization of a representative Norn cell niche in Xenium data, showing Norn cells (red) in close proximity to adventitial fibroblasts (cyan) and endothelial cells of the afferent- efferent arterioles (yellow). g) Heatmap of gene expression dynamics along pseudotime (Palantir), modeling the transition from adventitial fibroblasts to Norn cells. h) NicheNet ligand–receptor analysis identifying candidate signaling interactions with Norn cells. Dot plot shows ligand expression across sender populations. i) Using the precomputed hormone receptors from the Hormone cell atlas^64^ hormone production scores were calculated independently for the HKCA-cell and HKCA-nucleus datasets and subsequently integrated across both modalities to identify kidney and cell type specific endocrine signals for Norn cells. j) UMAP of LEPR expression across stromal populations in the HKCA-cell.

**Figure 4.**
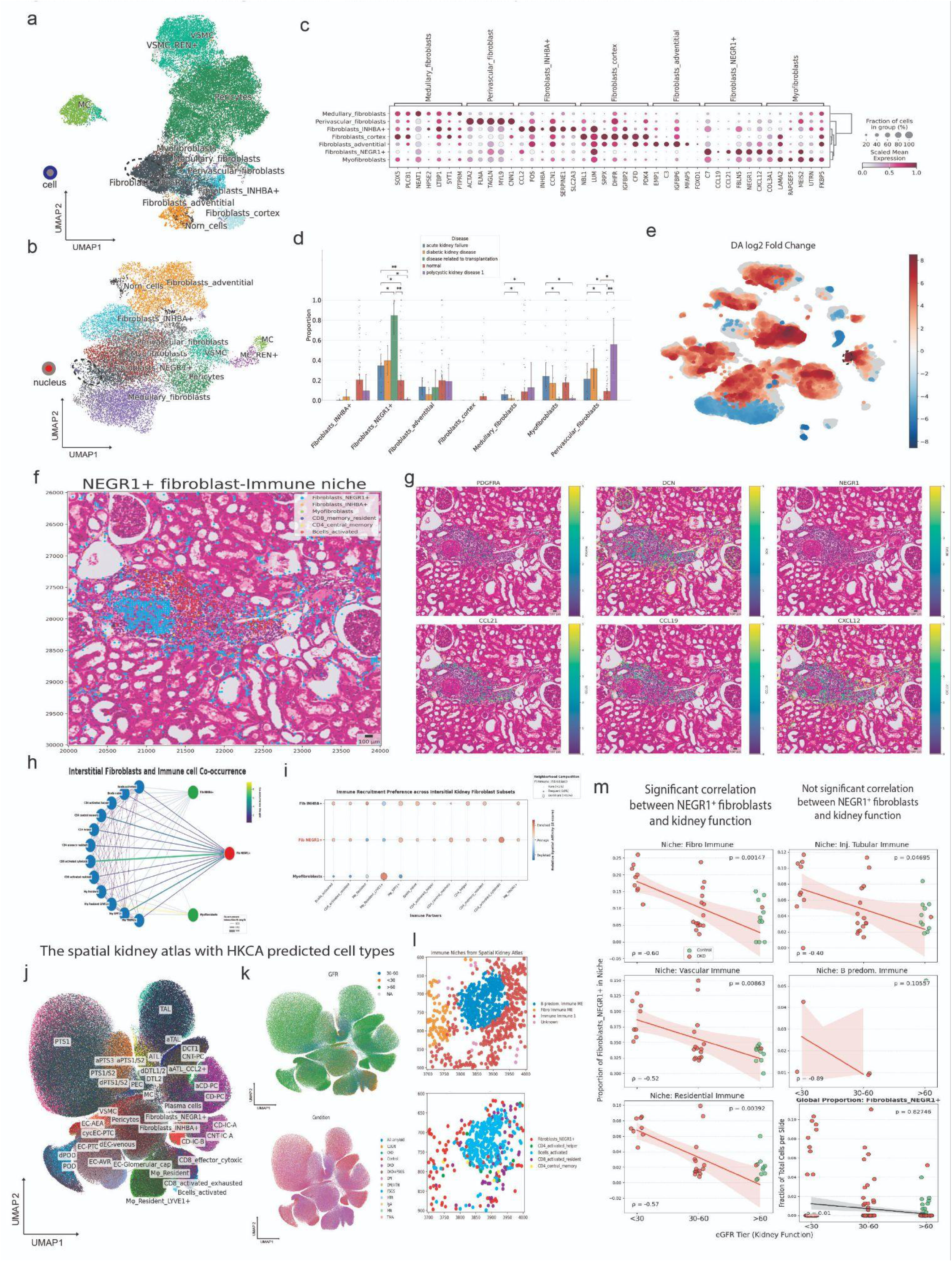
Immune attracting fibroblasts maintained in kidney homeostasis and are enriched in disease states. a,b) UMAP of stromal cells from the HKCA highlighting NEGR1 fibroblasts across HKCA-cell (part a) and HKCA-nucleus (part b) datasets (dashed circle). c) Dot plot of fibroblast marker genes, showing chemokine expression in distinct subsets, including CCL19/CCL21 in NEGR1 fibroblasts and CCL2 in INHBA fibroblasts. d) Bar plots of fibroblast subpopulation abundances across disease conditions. Proportions were calculated per sample and aggregated by condition; bars indicate mean ± 95% CI, with points representing individual samples. Statistical differences were assessed using Kruskal–Wallis tests with Benjamini–Hochberg correction (**P < 0.01). e) Spatial differential abundance analysis (Milo) comparing kidney transplant and normal tissue. Cells are colored by log-fold change in neighborhood abundance, NEGR1 fibroblasts shown in dashed circle. f) Spatial mapping of HKCA fibroblasts and immune cell types in VisiumHD tissue using a classifier trained on the HKCA. g) Spatial expression of marker genes for NEGR1^+^ fibroblasts (PDGFRA, DCN, NEGR1) and chemokines (CCL19, CCL21, CXCL12). h) Network representation of spatial co-occurrence as estimated by squidpy (radius=50-100um) between HKCA interstitial fibroblasts and immune cell types. i) Dot plot of spatial affinity between fibroblast subsets and immune cells. Dot size reflects co-occurrence frequency (radius=50-100um) and color indicates relative enrichment (Z-score). j) UMAP with the mapping of HKCA major cell types to the integrated kidney spatial atlas^8^ as predicted by a logistic regression model trained on the HKCA. k) UMAP with the eGFR function grouping (up) and the diseases included in the kidney spatial atlas. l) Example of a diabetic kidney disease sample from the spatial atlas showing the B cell predominant niche (Blue), the Fibroblast-Immune Niche (Orange) (top panel) and the associated predicted cell types (bottom panel) from the HKCA (NEGR1^+^ fibroblasts, red; CD4 T cells, green; CD8 T cells, purple; and B cells, cyan.) m) NEGR1^+^ fibroblasts mapped from the HKCA to the spatial atlas immune-associated niches. Healthy control samples (n=12) are depicted in green, while Diabetic Kidney Disease (DKD) samples (n=28) are shown in red. A linear regression trendline is fitted to the DKD cohort eGFR groups within each niche and the global spatial atlas. The bottom-right panel displays the global proportion of NEGR1^+^ fibroblasts across the entire spatial atlas. Spearman correlation coefficients (ρ) and corresponding p-values for the DKD cohort are annotated within each panel.

To minimize variability introduced by heterogeneous computational preprocessing, we acquired and remapped the raw FASTQ files using a unified workflow (see Methods) including same-reference reads alignment using STARsolo^31^ (including intronic reads to unify scRNA-seq and snRNA-seq), cell calling using EmptyDrops^32^, and ambient RNA correction using CellBender^33^ (Fig. 1a).

We next benchmarked alternative integration strategies using a curated subset of 10 datasets^6,7,9,11,12,22,23,25,27,34^ (Ext. Data Table 1) with harmonized reference annotations based on the original annotations by the data contributors. We compared label-independent and label-aware methods, including Harmony^35^, BBKNN^36^, Scanorama^37^, scVI^38^, scANVI^39^, scPoli^40^ and scGen^41^, across multiple feature-selection strategies.^42,43^ To avoid circularity between training labels and evaluation labels, prioritized label-unaware integration for production of the reference atlas. **scVI**^38^ provided the strongest performance (total score = 0.56) among label-unaware methods balancing batch correction and biological variance preservation and was thus selected as the primary atlas integration method (Fig. 1b, Ext. Data Fig. 4).

Joint integration revealed strong modality-dependent capture bias between single-cell and single-nucleus datasets. Immune cells showed greater abundance and resolution by scRNA-seq, whereas multiple epithelial populations, adipocytes and neurons are better captured by snRNA-seq (Fig. 1c, Ext Data Fig. 3a, 6). These differences likely reflect dissociation sensitivity and modality-specific recovery of cell types.

We further leveraged our HKCA to systematically evaluate modality-specific differential gene expression (Ext. Data Table 2) which is potentially generalizable for interpreting modality-based biases. snRNA-seq preferentially detected transcripts encoded by exceptionally large genomic loci, including ***MAGI2*** (∼1.4 Mb), ***DPP6*** (∼1.3 Mb), and ***WWOX*** (∼1.1 Mb), consistent with prolonged transcription and nuclear retention of incompletely processed pre-mRNAs. In contrast, scRNA-seq preferentially recovered highly abundant cytoplasmic transcripts, including ***GAPDH***, ***EEF1A1***, ***FTH1***, and ***FTL***, reflecting mature mRNAs that are rapidly exported from the nucleus and enriched in the cytoplasmic compartment Ext. Data Fig. 6c, 7). Based on these features, we developed a lightweight linear classifier named CellOrNuc to automatically infer the modality (cell or nucleus) with 100% accuracy at library level. Unexpectedly, at cell level, single-cell and single-nucleus profiles did not form two completely distinct populations. Instead, a small subset (4.3%) of single-cell profiles were classified as nuclei, hereafter referred to as **nucleus-like cells**, potentially representing stressed or damaged cells with reduced cytoplasmic RNA content. Conversely, a subset of single-nucleus profiles (0.45%) were classified as cells, hereafter referred to as **cell-like nuclei**, possibly reflecting nuclei that retained residual cytoplasmic RNA during nucleus isolation. The intermixing of single-cell and single-nucleus profiles is highly dependent on cell types, with epithelial and endothelial cells having the highest degrees of nucleus-like cells and cell-like nuclei (Ext. Data Fig. 7)

Rather than forcing the two modalities into a single overcorrected embedding, we retained the atlas as two linked components: **HKCA-cell** (816,020 cells) and **HKCA-nucleus** (215,308 nuclei) (Fig. 1d). We then performed the same clustering and annotation for HKCA-cell and HKCA-nucleus separately, identifying and removing doublet (108,777 cells and 51,674 nuclei) and low-quality or high-mitochondria droplets (433,609 cells and 62,823 nuclei), and flagging non-kidney contamination, including pancreatic acinar cells (43 cells) from one donor. These efforts resulted in 9 compartments, 63 cell types and 120 cell states with concordant marker gene signatures across the two modalities (Fig. 1e, Ext. Data Table 3, 4). Rare cell types and disease-associated cell states capturing subtle populations such as human kidney Schwann cells, ciliated cells, immune-attracting fibroblasts, disease-related podocytes and loop of Henle cells, a spectrum of renin and erythropoietin producing stromal cells, and adipocytes for the first time.

Together, this strategy produced a modality-aware human kidney reference of unprecedented resolution that preserves the complementary strengths of single-cell and single-nucleus RNA-seq while providing a unified annotation framework across datasets (Ext. Fig. 8, 9, 10, 11). The scale of the atlas enabled robust detection of major kidney lineages, rare, specialized populations and disease-associated cell states that were inconsistently resolved in individual datasets.

### Renin is produced by two mural cell populations

Among the 9 compartments, we first assessed the cell-state definitions within the stromal and mural compartments.^44,45^ Major stromal populations, including fibroblasts, mesangial cells, pericytes, and vascular smooth muscle cells, were readily recovered across datasets. However, the increased sampling depth and complementary strengths of single-cell and single-nucleus profiling enabled improved resolution of specialized mural populations involved in renin production. Renin production is a central component of kidney endocrine function and has traditionally been associated with juxtaglomerular and mesangial-like cells, including in recent reference atlases.^16^ In contrast, developmental and lineage-tracing studies have suggested that renin-producing cells arise from a broader mural-cell lineage that includes mesangial cells, vascular smooth muscle cells, and pericytes.^46,47^

We detected the expression of renin (*REN*) in vascular smooth muscle cells, mesangial cells and pericytes. Specifically, HKCA-nucleus resolved a mesangial REN-expressing population (MC-REN^+^) (Fig. 2a), while HKCA-cell identified a vascular smooth muscle REN-expressing population (VSMC-REN^+^) (Fig. 2b) that co-expressed canonical contractile markers including *ACTA2*, *MYH11*, and *TAGLN* (Fig. 2c). Transcriptional harmonization between MC-REN^+^ and VSMC-REN^+^ cells revealed a shared core program, including *MECOM*, *HPSE2*, *PLCL1* and *PDE10A*^46^ (Ext. Data Fig. 10), supporting a broader mural-cell lineage program.^48,49,47,50,51^ Additionally, these findings suggest that previous human kidney atlases may have incompletely captured the full diversity of renin-producing cells because of modality-specific sampling biases.

To define their anatomical context, we projected atlas-derived gene expression signatures onto a published VisiumHD spatial transcriptomic data from a healthy human kidney.^52^ Spatial mapping localized MC-REN^+^ and VSMC-REN^+^ cells to both mesangial and perivascular compartments (Fig. 2d, e), supporting the existence of anatomically distinct renin-producing populations in the adult human kidney. To investigate whether these anatomically distinct REN^+^ populations are also subject to different modes of physiological regulation, we examined the expression of receptors implicated in renin control, including adrenergic (*ADRB1*)^53^, prostaglandin (*PTGER4*)^54^ and angiotensin (*AGTR1*)^55^ receptors in these renin producing populations (Fig. 2c). VSMC-REN^+^ cells preferentially expressed receptors associated with sympathetic and prorenin signaling, including *ATP6AP2* (FC=+11.24, padj<0.001), *ADRB1* (FC=+4.35, padj=0.5) and *ADRA1*(FC=+4.79, padj<0.001) compared to MC-REN^+^ and weak response to prostaglandins. In contrast, the MC-REN^+^ cells preferentially expressed receptors associated with **feedback regulation of the renin–angiotensin–aldosterone axis**, including the angiotensin II receptor *AGTR1* (logFC=-3.0, padj<0.001) and the aldosterone responsive receptor *NR3C2* (logFC=-8, padj<0.001). These differences suggest that VSMC-REN^+^ may directly respond to neuronal and local prorenin levels at the vascular bed compared to MC-REN^+^ which respond to the feed-forward loop of the renin-angiotensin-aldosterone system.

Together, these findings provide evidence that renin production in humans is distributed across two mural cell types, extending observations from developmental studies and refining our understanding of the cellular organization of the renin-producing compartment.^47^

### A perivascular stromal niche supports erythropoietin-associated Norn cells

In addition to renin production, the kidney serves as the primary source of systemic erythropoietin (EPO), a hormone that stimulates erythropoiesis and whose production is modulated by tissue oxygen availability. Recent studies identified Norn cells as a rare stromal population responsible for EPO production. ^18,56^ The increased stromal cell and sample numbers of the HKCA enabled robust identification of Norn cells as a rare subtype of adventitial fibroblasts, which expressed previously described gene markers including *CXCL14, TCF21, PRRX1*^56^. However, the HKCA harmonization process also revealed that the Norn cell state is characterized by the expression of suspension unbiased markers, including *IGFBP6, BNC2, SFRP4* (Fig. 3a,b,c Ext. Data Fig. 10a,b).^57,58^

To define the spatial context of Norn cells, we mapped HKCA annotations onto a Xenium spatial transcriptomics dataset of a tumor-adjacent healthy kidney tissue.^59^ Norn cells localized to a perivascular niche adjacent to afferent and efferent arterioles and were consistently found in proximity to adventitial fibroblasts and resident macrophages (Fig. 3d–f), suggesting that local cellular interactions may contribute to emergence and maintenance of Norn-cell identity. Along the continuum from adventitial fibroblasts to Norn cells, we identified a progressive activation of genes including *SOD3*, *PPRX1*, *IGFBP6*, and *LEPR*, preceding establishment of the mature Norn transcriptional identity (Fig. 3g). Additionally, the receiver-centered ligand– receptor analysis^60^ identified signaling pathways associated with the stromal cell niche (Fig. 3h), such as PDGF ligands derived from adventitial fibroblasts, extracellular-matrix-associated signaling through decorin (*DCN*), and a GAS6–AXL signaling axis associated with hypoxia-response and cell-survival programs.^61^ Norn cells also showed enriched expression of leptin receptor (*LEPR*), suggesting potential responsiveness to endocrine signals early in their developmental trajectory.^62,63^ To discover additional hormonal regulatory codes at a whole-body level, we leveraged the Hormone cell atlas^64^ and identified resistin and vaspin along with leptin to be amongst the top interacting hormones for Norn cells in a paracrine manner. (Fig. 3i, j, Ext. Data Fig. 14). These results suggest that adventitial fibroblasts acquire a specialized fate for EPO production controlled by endocrine and local signals.

It is possible therefore that kidney diseases which chronically alter this perivascular niche and induce hypoxia, such as vasculitis^65^ or CKD^66^ might impact the abundance of Norn cells. Indeed, we observed higher numbers of Norn cells in transplant recipients, consistent with the known effect of transplantation in improving CKD related anaemia^67^ (Ext. Fig. 12). Another chronic disease of the kidney which is known to present with both anemia and vascular damage is IgA nephropathy.^68,69^ Mapping HKCA annotations onto a publicly available Xenium spatial transcriptomics sample of IgA nephropathy kidney^70^, we found substantial remodeling of the Norn-cell microenvironment. Compared with healthy kidney tissue, Norn-associated neighborhoods showed reduced representation of adventitial fibroblasts and increased co-localization of two disease-associated INHBA^+^ and NEGR1^+^ fibroblast populations newly identified by our HKCA, together with activated resident CD8^+^ T cells and mast cells (Ext. Data Fig. 15). Collectively, these observations in combination with adventitial nature of Norn cells suggest that the remodeling of the stromal niche in CKD combined with disease specific vascular damage may directly affect Norn cells and EPO production.

### Disease-associated fibroblast states

Since the HKCA was compiled as a compendium of both healthy and diseased samples, we next performed a systematic survey of disease-associated cell states using Milo^71^ (Ext. Fig. 12, 13). We have identified disease specific cellular abundance (spatialFDR <0.05) in aPTS3, NEGR1^+^ fibroblasts and SPP1^+^ macrophages in AKI, while ADPKD was enriched for aPTS1/S2, INHBA^+^ fibroblasts, adventitial fibroblasts, perivascular fibroblasts and monocytes, and transplant samples showed an increase in neighborhoods of aPTS3, aATL-CCL2^+^, aCD-PC, classical monocytes, damaged EC-venous and NEGR1^+^ fibroblasts. Stemming from this analysis, we first investigated adventitial, medullary fibroblasts and a spectrum of transcriptionally distinct interstitial fibroblast populations with highly concordant transcriptional signatures in the HKCA-cell and HKCA-nucleus (Fig. 4a,b).

The newly identified INHBA^+^ fibroblast population displayed an activated remodeling program characterized by high expression of *INHBA, CCL2, SERPINE1, CCN1*, and *CCN2*, consistent with extracellular matrix remodeling and Activin-associated signaling (Fig. 4c, Ext. Data Fig. 10). Activin-driven stromal programs have been described in other organs including the kidney, where Activin A signaling contributes to fibroblast activation and remodeling of the local tissue microenvironment in cancer.^72,73^ The differential abundance analysis (Ext Data Fig. 12, 13) revealed that INHBA^+^ fibroblasts were expanded in the adult polycystic kidney disease cohort (n=8), suggesting that the TGF-beta/SMAD pathway might be associated with this fibroblast population during this disease process.^72^

In contrast to activin’s role in fibroblast remodeling, NEGR1 is a neurite enriched protein of the immunoglobulin superfamily implicated in fatty acid regulation and fibroblast growth factor stabilization in fibroblast progenitors and pre-adipocytes.^74,75^ Through GWAS analysis it has been implicated with worsening eGFR in the UK biobank cohort ^76^ but lacking kidney cell type association. Through an iterative annotation process the HKCA identified a NEGR1^+^ expressing fibroblast population with a striking enrichment in kidney diseases especially transplant rejection and acute kidney injury (Fig. 4d, e; Ext. Data Fig. 7). Although present under homeostatic conditions, NEGR1^+^ fibroblasts in the HKCA-cell are exceptionally abundant in transplanted kidneys undergoing rejection (Fig. 4d, e). Concurrently, they selectively expressed the homeostatic chemokines *CCL19*, *CCL21*, and *CXCL12*, which resemble the stromal populations described across lymphoid and peripheral tissues that regulate immune-cell trafficking and maintenance ^77,78^.

HKCA-guided spatial mapping onto VisiumHD from healthy donor localized NEGR1^+^ fibroblasts to interstitial regions adjacent to dense lymphoid infiltrates (Fig. 4f, g). Neighborhood analysis amongst the interstitial fibroblast subsets further demonstrated preferential association with innate and adaptive immune populations, including resident macrophages, memory CD4^+^ T cells and B cells, supporting further the specialized role of NEGR1^+^ fibroblasts in organizing local adaptive immune responses (Fig. 4h, i).

To evaluate the clinical relevance of this stromal niche, we next projected HKCA fibroblast annotations onto an independent published spatial transcriptomic atlas that includes samples of diabetic kidney disease (DKD).^8^ HKCA-guided reannotation successfully recovered the same immune-associated NEGR1^+^ fibroblast neighborhoods (Fig. 4j–l). Notably, the proportion of NEGR1^+^ fibroblasts in immune-associated niches (fibroblast-immune, vascular-immune and residential immune) was significantly inversely correlated with kidney function (determined by estimated glomerular filtration rate (eGFR)) in DKD (n=28) vs healthy controls (n=12) (ρ = −0.60, p = 0.0015), but not in all immune niches (Fig. 4m). Therefore, providing further support for the role of this interstitial population as central in organising specific immune-stroma niches in the kidney and its relevance to kidney function in DKD.

### A Loop of Henle population linked to genetic risk for reduced kidney function

In addition to the aforementioned fibroblasts, epithelial cell state subsets were also captured in our global disease association analysis. aPTS1/S2 state was enriched in autosomal dominant polycystic kidney, while aPTS3 was more prominent in transplant kidney disease. Additionally, a distinct CCL2^+^ ascending thin limb (aATL_CCL2^+^) cell population emerged as one of the most prominent injury-associated cell states in kidney transplant. (Fig. 5a) CCL2 is a well-established mediator and biomarker of kidney injury and allograft inflammation, including monocyte/macrophage recruitment during rejection and association of urinary CCL2 with long-term renal allograft outcome.^13,79^

**Figure 5.**
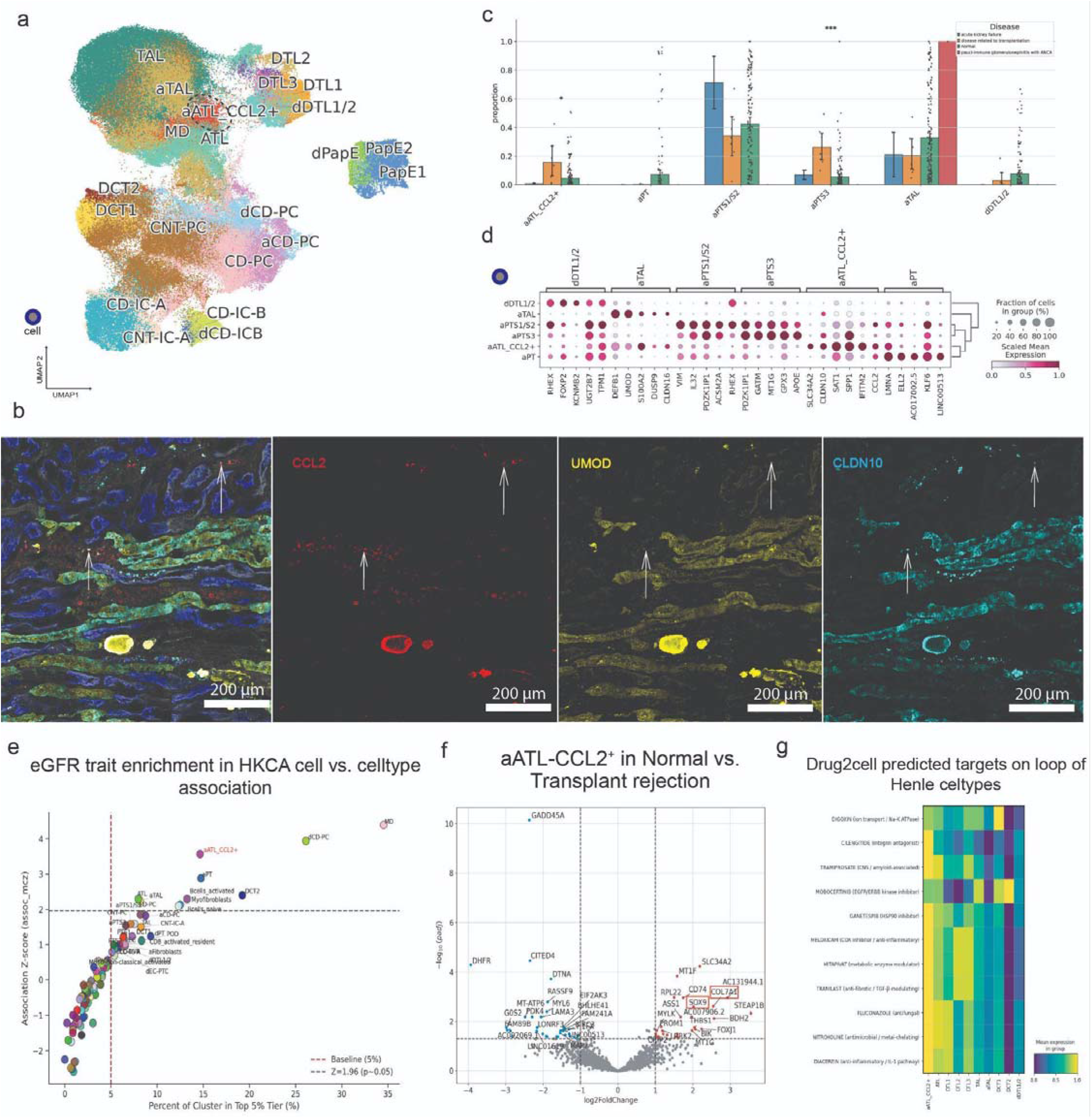
A Loop of Henle epithelial population linked to genetic risk for reduced kidney function. a) UMAP of epithelial cells from the HKCA-cell highlighting the CCL2-expressing ascending thin limb Loop of Henle population (black dashed circle). b) Dot plot showing marker gene expression for disease-associated Loop of Henle cell types and proximal tubular epithelial cells. c) Bar plots of relative abundances of epithelial cell types across disease conditions in the HKCA-cell. Proportions were calculated per sample and aggregated by disease; bars indicate mean ± 95% CI, with points representing individual samples. Statistical differences were assessed using Kruskal–Wallis tests (≥3 samples per disease) with Benjamini– Hochberg correction (*P < 0.05, **P < 0.01, ***P < 0.001). d) Representative immunohistochemistry (IHC) images showing co-localization of CCL2 with UMOD (thick ascending limb) and CLDN10 (thin ascending limb), confirming the aATL-CCL2 population (n=1). White arrows indicate CCL2 epithelial cells within the ascending limb (scale bar, 200 μm). e) Scatter plot showing GWAS enrichment for creatinine-based eGFR across cell types. The x-axis represents the proportion of cells in the top 5% of scDRS scores, and the y-axis shows association Z-scores (dashed lines indicate thresholds). f) Volcano plot of differential gene expression between kidney transplant disease and normal tissue within the aATL-CCL2 population (pseudobulk analysis). Significant genes (padj < 0.05, |log FC| > 1) are shown in red. g) Heatmap of Drug2Cell enrichment scores across Loop of Henle cell types.

Immunohistochemical analysis of cadaveric kidney glomeruli (Fig. 5b) validated this previously unrecognized epithelial state by identifying CCL2^+^CLDN10^high^ epithelial cells within the *UMOD*^-^ ascending limb, confirming the presence of aATL-CCL2^+^ cells in human kidney tissue. Unlike canonical ascending thin limb cells, aATL-CCL2^+^ cells were markedly enriched in transplant rejection (n=6) while remaining infrequent in healthy kidneys (n=47) (Fig. 5c, Ext. Data Fig. 12). The aATL-CCL2^+^ population expresses a coordinated injury-remodeling signature marked by *CCL2, IFITM2, SPP1, SAT1* and *WFDC2*, ^80,81,82,83^ suggesting chemokine signaling, epithelial stress responses, interferon response and metabolic dysfunction within a single epithelial cell state. (Fig. 5d, Ext. Data Fig. 9)

We next asked whether disease-associated epithelial states captured by the HKCA also contribute to inherited variation in kidney function. We integrated HKCA annotations with GWAS summary statistics for creatinine-based eGFR kidney function (eGFRcrea)^84^ using MAGMA gene-set analysis,^85^ and refined this signal by applying single-cell disease relevance scoring (scDRS).^86^ scDRS localized polygenic kidney-function risk to aATL-CCL2^+^ cells compared with neighboring Loop of Henle populations, along with damage principal cells and macula densa cell types known to control the kidney’s filtration function. (Fig. 5e, Ext. Data table. 5) Together, these results identify aATL-CCL2+ cells as a site of convergence between an injury-associated epithelial state and the expression of genes implicated by inherited variation in kidney function. (Ext. Fig. 16)

Finally, to identify possible therapeutic targets associated with the aATL-CCL2^+^ transcriptional program, we performed Drug2Cell^87^ analysis across Loop of Henle epithelial cells (Fig. 5g). Whereas canonical Loop of Henle cells were primarily enriched for ion transport–related pathways, aATL-CCL2^+^ cells additionally showed selective targeting by various drugs, including the integrin signaling inhibitor cilengitide,^88^ and the anti-fibrotic agent tranilast.^89^ This analysis suggests that disease associated, and genetic risk-linked aATL-CCL2^+^ population may be potentially targetable by existing drugs in treating functional decline related to fibrosis.

### An injury-associated podocyte state

While injury-associated tubular populations have been widely described, much less is known about the diversity of states within human podocytes. The increased sampling depth of the HKCA resolved three reproducible podocyte states after gene harmonization (Fig. 6a, b). In addition to a canonical homeostatic population (AIF1^−^ POD), we identified a distinct AIF1^+^ POD_ECM state characterized by high expression of *AIF1*, *BCAM*, *SPOCK2*, *CXCL14*, and *FGF1*, together with extracellular matrix-associated genes (Fig. 6c).^90–92^ A third population (dPOD) identified in the HKCA-cell exhibited elevated expression of AP-1 family transcription factors and a large fraction represented nucleus-like cells, possibly due to an acute stress-response program (Fig. 6d),^93,94,95,96^ or dissociation during library preparation.

**Figure 6.**
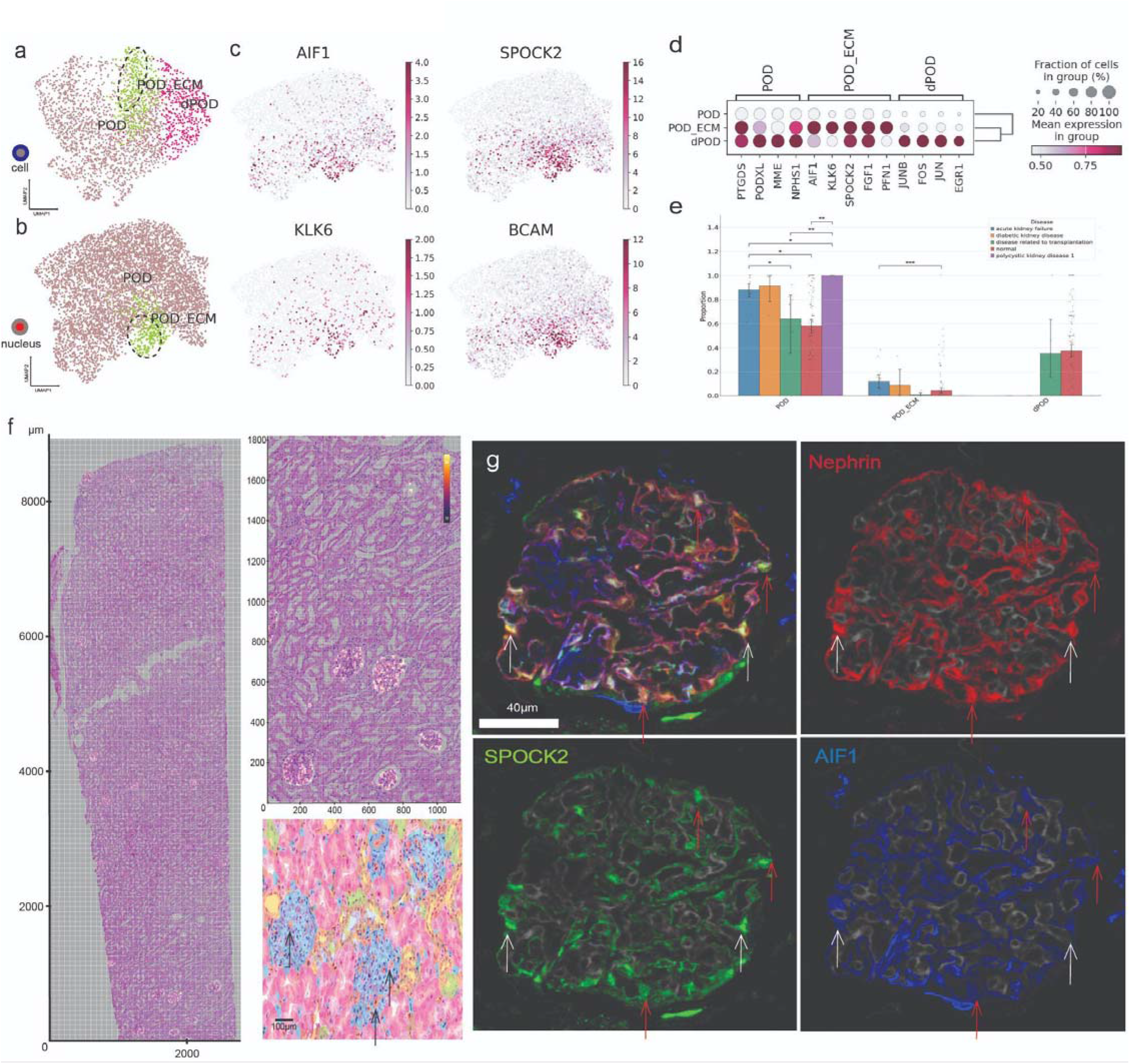
Podocyte populations in kidney. a,b) UMAP embeddings of podocyte populations from HKCA-cell and HKCA-nucleus data, highlighting a distinct ECM-modulating subset (POD_ECM) that segregates from canonical podocytes (POD). c) UMAPs showing expression of key marker genes (AIF1, SPOCK2, KLK6, BCAM). d) Dot plot of marker gene expression across podocyte populations. e) Quantification of podocyte subpopulation proportion from the HKCA, across disease conditions. Bars show mean ± 95% CI. g) Immunofluorescence validation in glomeruli of cadaveric kidney showing co-localization of nephrin (NPHS1) with SPOCK2 and AIF1, confirming the presence of the ECM_POD population (white arrows indicate AIF1^+^ podocytes, red arrows indicate AIF1^-^ podocytes). f) Spatial transcriptomics (Xenium) of a healthy nephrectomy kidney (n=1)^59^ showing glomerular localization of AIF1 expression and mapping of POD-ECM cells (orange / black arrows) based on the combined HKCA-cell and HKCA-nucleus derived predictions.

We next asked whether these transcriptional states were associated with kidney disease. Among the three populations, POD_ECM showed strong disease association, being reproducibly detected in both the HKCA-cell and HKCA-nucleus atlases and significantly enriched in acute kidney failure compared with healthy samples (Fig. 6e). To determine whether this state could be localized within a healthy tissue, we projected HKCA podocyte annotations onto the published Xenium (10X Genomics).^59^ POD_ECM cells were mapped to glomeruli, where AIF1 expression was selectively enriched (Fig. 6g). Immunofluorescence staining of cadaveric human kidney glomeruli further demonstrated the coexistence of AIF1^+^ and AIF1^−^ nephrin-positive (*NPHS1*) podocytes within individual glomeruli, raising the possibility that these represent dynamic podocyte states associated with injury, or constitutional repair of the glomeruli (Fig. 6f). Therefore, the harmonized analysis of the HKCA-cell and HKCA-nucleus resolved POD_ECM as a responsive podocyte state specific for extracellular matrix remodeling in the glomeruli and enriched in acute kidney disease.

### HKCA-guided urine cell profiling enables kidney disease stratification

To demonstrate the utility of the HKCA for automated annotation of single cell transcriptomics data, we trained a cell type-prediction model (scArches) based on the HKCA^97^, to automatically analyze publicly available urine-derived single-cell transcriptomic data (83,035 cells) from diseased samples including acute kidney injury (n=35), focal segmental glomerulosclerosis (n=23), kidney transplant recipients (n=1), membranous nephropathy (n=3), lupus nephritis (n=1) and those with arterionephrosclerosis (n=2).^98–101^. This transfer learning approach yielded high-confidence cell-state labels across epithelial, immune, and stromal compartments and across independent batches (Fig. 7a). The automatically generated labels are strongly concordant with the cell state annotations given by the original publications showing that the HKCA is a robust framework for interpreting independent urine-derived datasets. In addition, the HKCA was able to identify NK cells—absent from original annotations—and show specific enrichment in the membranous nephropathy urine samples (Fig. 7b, c).

**Figure 7.**
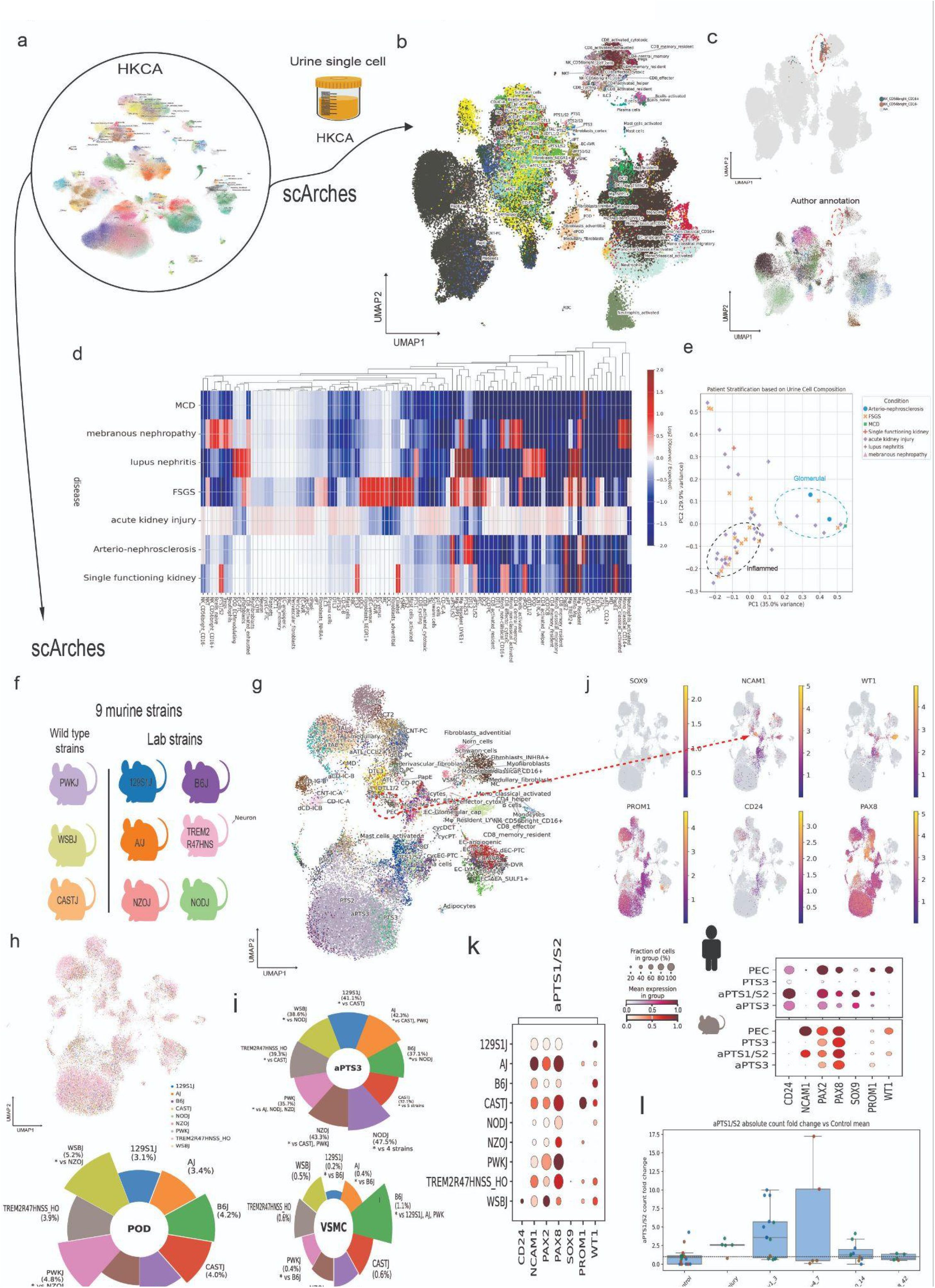
The HKCA can predict cell states in urine samples and mice. a, b) Reference mapping of urine-derived single-cell transcriptomes^98–101^ onto the HKCA using scArches. Cells from diseased samples acute kidney injury (n=35), focal segmental glomerulosclerosis (n=23), kidney transplant (n=1), membranous nephropathy (n=3), lupus nephritis (n=1) and those with arterionephrosclerosis (n=2) are projected into the latent space and colored by predicted cell type, showing alignment with major renal epithelial, stromal, and immune populations. c) Predicted annotations show high concordance with original dataset labels, with high confidence across most cells and new cells identified by the reference mapping. d) Heatmap of cell type enrichment across disease conditions based on scArches predictions. Values represent log observed-to-expected proportions, highlighting disease-associated shifts in epithelial and immune composition. e) Principal component analysis (PCA) of patient-level cell type composition, showing separation of the urine samples by disease. f) Reference mapping of IGVF consortium dataset across 9 mouse strains’ kidneys sequenced by SPLiT-seq in a multiplexed cross-tissue experiment.^103^ g) Predicted labels by the HKCA using the scArches embedding on the IGVF dataset^103^. h) UMAP showing the strain contribution from the IGVF dataset^103^ to each individual cluster of predicted labels. i) Proportion of top 3 significantly different cell types across strains (Ext. Data Fig. 17) j) UMAPs showing the expression of renal progenitor markers (*CD24, PROM1, SOX9, WT1, PAX2, PAX8*) in the *Ncam1^+^*epithelial cell population. k) Cross-species mapping of the progenitor markers in the predicted cell types. The expression of renal progenitor markers (*CD24, PROM1, SOX9, WT1, NCAM1*) across 9 mouse strains in the predicted aPTS1/S2 cell population. l) Abundance of aPTS1/S2 cells in 4 single cell datasets of ischemic mouse kidney injury during the initial 7 days post injury. The dashed line shows control level.

Systematic evaluation of urine-derived cellular composition revealed disease-specific enrichment patterns that reflected the underlying clinical pathobiology (Fig. 7d). For example, inflammatory conditions, such as lupus nephritis, were characterized by an influx of activated myeloid and lymphoid populations in the urine, whereas membranous nephropathy displayed increased NK-cell representation. In contrast, samples from patients with chronic kidney injury, including kidney transplant recipients and those with arterionephrosclerosis, showed enriched proportions of injured and segment-specific epithelial states, particularly from the proximal tubule and thick ascending limb (TAL). Urine samples from patients with focal segmental glomerulosclerosis (FSGS) displayed a heterogeneous immune and epithelial profile, mirroring its diverse clinical presentation. ^34,102^

Principal component analysis (PCA) of these cellular proportions demonstrated distinct clustering by disease category, with clear separation between inflammatory and chronic injury-associated states (Fig. 7e). While these conditions formed relatively cohesive groups, acute kidney injury samples showed greater variation, likely reflecting the diverse etiologies and temporal stages of injury captured within the urinary compartment.

### HKCA-guided mouse profiling enables cross-strain and human comparisons

We further extrapolated the transfer learning approach to evaluate cross-species differences, using the Impact of Genomic Variation on Function (IGVF) Consortium’s dataset containing mouse strains commonly used in research analyzed using SPLiT-seq.^103^ (Fig. 7f, g) The label transfer across 9 mouse strains, including wild type and inbred strains (Fig. 7h), automatically recovered high quality cell annotations, largely consistent with the labels provided by the original authors. We identified aPTS3 as the most prominent cell state (Fig. 7i, Ext. Fig. 17), indicating that the transcriptional program defining human aPTS3 represents a conserved physiological state, marked by the expression of genes related to oxidative stress and epithelial repair, which appears constitutively active within the proximal tubular cells of healthy mice.

Interestingly, cross-species label transfer using scArches resolved the mouse *NCAM1* epithelial cell population into two human-derived cell states: parietal epithelial cells (PECs) and adaptive proximal tubule S1/S2 (aPTS1/S2) (Fig. 7j).^104^ Notably, *NCAM1* and *WT1* were primarily restricted to the PEC compartment within HKCA-cell and HKCA-nucleus, with negligible expression in proximal tubule populations. In contrast, established human renal progenitor-associated markers, including *CD24*, *PROM1* (*CD133*), and *SOX9*, were specifically enriched in the aPTS1/S2 population.^105,106^ Cross-strain analysis revealed that this transcriptional signature was uniquely recapitulated in the wild-type WSBJ strain (Fig. 7k), suggesting strain-specific divergence in the adaptive proximal tubule cell program. Finally, we replicated the label mapping analysis in 4 independent kidney mouse injury datasets ^107–110^ to further evaluate proximal tubule responses to injury. We consistently found aPTS1/S2 cell number increases during the progression of ischaemic injury as modeled by experimental artery ligation to the kidney suggesting this population is injury-responsive. (Fig. 7l)

## Discussion

In this study, we present the HKCA, a systematically integrated reference atlas spanning independently generated human kidney sc/snRNA-seq datasets from across multiple studies, institutions and disease contexts (Ext. Fig. 2). We have not only unified the cell state labels across studies but have demonstrated the power of large-scale data integration by identifying previously unknown cell states. The HKCA demonstrates robust cross-modality concordance at the level of cell identity and transcriptional programs,^18^ and a harmonized annotation (Fig. 1 and Ext. Fig. 8, 9, 10, 11) revealed a rich landscape of 120 kidney cell states, including disease-associated populations, with consistent marker signatures across modalities.

Using the HKCA, we identified human EPO-producing Norn cells ^56^ that were reported in large foundation model-based analyses of scRNA-seq data^18,111^. We found that Norn cells are located with a distinct perivascular niche that exhibits dynamic remodeling in the IgA nephropathy micro-niche. (Fig. 3, Ext. Fig. 21) These analyses highlight the importance of stromal–vascular interactions in maintaining Norn cell identity and suggest that disruption of this niche is associated with disease. Notably, Norn cells share transcriptional features (*CXCL14*, *LEPR*) with previously described leptin-responsive stromal preadipocyte populations, suggesting a broader role of leptin as a stromal cellular growth factor, as shown by alignment with the Hormone Cell Atlas.^64,75^

Our analysis also revealed that renin expression is distributed across both mesangial and vascular smooth muscle compartments, rather than a single cell type as previously proposed. This is consistent with prior developmental studies demonstrating that renin-lineage cells arise from FOXD1 stromal progenitors^47,112^, which give rise to mesangial cells, vascular smooth muscle cells, and pericytes. In this context, the shared transcriptional program observed between MC-REN and VSMC-REN populations (Fig. 2) may reflect a conserved lineage identity, with divergence driven by anatomical localization and microenvironmental cues. These findings support a model in which vascular smooth muscle cells can activate renin expression under physiological or stress conditions.^46^ More broadly, this distributed organization of renin production may provide functional flexibility in maintaining renal hemodynamics, allowing multiple mural cell populations to contribute to the regulation of the renin–angiotensin system.

A key strength of the HKCA lies in its ability to resolve disease-relevant cellular states and link them to genetic susceptibility. We identify an adaptive loop of Henle epithelial population (aATL-CCL2, Fig. 6) that is enriched in chronic immune-mediated injury, associated with genetic risk for reduced kidney function, and undergoes a transcriptional shift indicative of maladaptive repair and fibrosis in transplant rejection. These findings connect genetic risk, cellular state transitions, and clinical pathology within a unified framework. Similarly, the identification of NEGR1 immune-attracting fibroblasts (Fig. 4) in both health and disease highlights a stromal cell population which organizes local immune–stromal interactions responsible for spatial immune organization described during kidney development and disease.^27,113^ The expanded kidney immune cell type identification provided by the HKCA spanning resident lymphoid subsets (effector memory CD4, CD8 and activated memory B cells) enabled us to further delineate the role of NEGR1^+^ in orchestrating local adaptive immunity as confirmed by spatial analysis. The alignment of the HKCA and the spatial atlas of diabetic kidney disease,^8^ further established the association of this cell state with kidney function decline in the diabetic kidney disease. Therefore, the HKCA establishes NEGR1^+^ fibroblasts as architects of immune niches specific to kidney function and demonstrates the power of high-resolution reference atlases to anchor and contextualize disease-state spatial transcriptomics and relay information on progressive kidney decline.

Beyond tissue-based analysis, our results demonstrate that the HKCA enables generalizable annotation and disease inference in independent human urine and mouse sc/snRNA-seq datasets. Notably, urine-derived samples originating from diseases not represented in the atlas could be accurately mapped and stratified based solely on inferred cellular composition. The ability to position non-represented diseases within this framework highlights that cell type composition encodes sufficient biological signal to resolve disease states independently of prior inclusion in the reference, suggesting cellular and transcriptional embeddings can be used in disease classification.^114^ Similarly, the HKCA can act as a benchmarked, modality aware reference to enable cross-species modeling of kidney function. Our analysis showed that both the mouse strain and cross-species transcriptional variability should be taken into consideration when interpreting the functional relevance of cell states in healthy and in ischaemic disease states. (Fig. 7)

Several limitations should be considered. First, the inability to fully integrate single-cell and single-nucleus transcriptomic datasets reflects both technical differences in capturing modalities and underlying biological biases, such as transcript localization and dissociation sensitivity. To that extent, our CellOrNuc classifier provides a new approach for the community to quantify and potentially mitigate modality-specific batch effects as a data cleaning method.^115^ Second, the atlas aggregates datasets generated across diverse protocols, cohorts, and disease contexts, which introduces confounding variances that current data cleaning and data integration methods may not be able to handle; for example, disparate capture efficiencies often exist between sc/snRNA-seq, particularly for matrix-embedded populations like mesangial cells, adipocytes or fibroblasts. Similarly, there is a scarcity of samples for certain diseases such as ADPKD and immune-mediated glomerulonephritis which can hinder disease specific analysis and therefore the systemic interrogation of the HKCA can help design future cohorts of increased diversity. Third, urine-based analyses rely on cells shed into the urinary space, which may incompletely represent the tissue composition of the urinary tract and are likely biased toward specific epithelial and immune populations. Finally, while compositional analyses reveal strong associations with disease states, causal relationships between specific cell populations and disease progression should be investigated in the future under established time series specimen collection protocols.

Despite these limitations, the HKCA provides a uniquely comprehensive and scalable framework for studying human kidney biology and modeling kidney disease. More broadly, our findings illustrate how large-scale integration of independent single-cell datasets can transform fragmented transcriptomic studies into genetically and spatially informed reference frameworks. By linking cell states to inherited risk, tissue architecture and disease remodeling, the HKCA provides a foundation for mechanistic interpretation of human kidney disease and future precision nephrology.

## Methods

### Raw data processing and generation of count matrices

All raw sequencing data were reprocessed using the STARsolo pipeline (STAR v3.0.0)^31^ as implemented by the CellGeni group (https://github.com/cellgeni/STARsolo). This approach enabled standardized preprocessing across datasets, including those generated by collaborating groups, while maintaining data governance requirements. Briefly, FASTQ files were aligned to the human reference genome (GRCh38-2020-A) including intronic sequences, and gene–cell count matrices were generated using default STARsolo parameters optimized for droplet-based single-cell RNA sequencing. To distinguish true cells from background barcodes, empty droplets were identified and removed using the EmptyDrops^32^ method. This step ensured the retention of high-confidence cell-containing droplets while excluding ambient RNA contamination. We subsequently evaluated for reprocessing replicate droplets by assessing barcode clashing as implemented in Scanpyplus.^43^ Finally, all matrices underwent Cellbender v0.2^33^ for ambient RNA correction per dataset and the corrected matrices were used as the raw matrices for subsequent analysis.

### Dataset selection and benchmarking cohort construction

For benchmarking, 10 datasets were manually annotated to generate ground-truth labels for cell-type-aware and cell-type-unaware evaluation (Ext. Data Table 1). These labels were curated independently prior to integration and used as the reference standard for benchmarking performance. We compared multiple integration methods, including Harmony^35^, Scanorama^37^, BBKNN^36^, scVI^38^, scANVI^39^, scPoli^40^, and scGen^41^, under multiple parameter settings. Benchmarking was performed using both combined and modality-separated integration strategies for single-cell and single-nucleus data, and across different highly variable gene (HVG) feature sets, including 5,000 HVGs, 15,000 HVGs, and a gene set derived by the Deeptree algorithm ^43^ from lineage-informative markers.

Performance was evaluated using scIB metrics as implemented by the HCA scAtlasTB toolbox^116^ to assess batch mixing and biological conservation. The best-performing label-unaware method was selected for downstream atlas construction based on its overall balance between integration quality and preservation of biologically meaningful structure. As a result, scVI^38^ with 15,786 genes identified by the Deeptree algorithm^43^ were used for integration. Raw count matrices were provided as input, with sample identity specified as a batch covariate. To account for technical variation, the proportions of mitochondrial, ribosomal, and hemoglobin-derived transcripts were included as continuous covariates.

The scVI model was initialized using a zero-inflated negative binomial (ZINB) likelihood with a two-layer neural network architecture comprising 256 hidden units per layer. Training was performed on a GPU for a maximum of 400 epochs using a 90:10 training-to-validation split and early stopping based on validation performance. The resulting latent representation was used for downstream visualization, clustering, and analyses.

### Cross-modality annotation and harmonization

Due to persistent modality-specific differences between single-cell and single-nucleus RNA sequencing data, including biases in transcript capture and gene detection, a unified integration of both modalities could not be achieved without compromising biological signal. Therefore, scRNA-seq and snRNA-seq datasets were processed and annotated as separate objects (HKCA-cell and HKCA-nucleus) based on the dataset of origin. Despite this, we found a partial integration from some specific clusters which has a contribution from both single-cell and single-nucleus. (Ext. Data Fig. 14)

To ensure consistency across modalities, Leiden algorithm as implemented in scanpy^117^ was used at initial clustering resolution of 1.0 for both objects and subsequently we applied a shared marker-based annotation framework using curated gene signatures derived from canonical kidney cell types and lineage-resolved programs (https://github.com/Peng-He-Lab/Kidney_integration). Annotation was performed using the Scanpyplus package^118^, which enabled systematic scoring of gene signatures and hierarchical assignment of cell identities. This approach allowed harmonization of cell type labels across modalities while preserving modality-specific transcriptional features, resulting in a consistent and biologically coherent annotation of epithelial, stromal, vascular, and immune populations across both atlases. Additional harmonization was performed for cross-modality detection similarities based on Cellhint^119^ and the final harmonized annotations were used for marker extraction.

Marker genes were identified using the Scanpyplus.filtered_markers framework applied to the scRNA-seq dataset. Differential expression analysis was performed using a Wilcoxon rank-sum test, and markers were filtered based on adjusted P value (< 0.05), minimum in-group expression fraction (> 0.25), minimum log fold-change (> 0.5), and maximum out-group expression fraction (< 0.75). Markers were ranked per cell type, and the top genes were selected for downstream visualization and annotation. Gene expression patterns were visualized using scaled dot plots to facilitate comparison across cell populations.

### Identification of modality-associated genes

To identify genes associated with single-cell (scRNA-seq) and single-nucleus (snRNA-seq) profiling, we combined cells from the HKCA-cell and HKCA-nucleus after removing mitochondrial and ribosomal genes. Gene expression values were library-size normalized and log-transformed [log1p(counts)]. Cells were labeled according to sequencing modality and used to train a Random Forest model (scikit-learn v1.7.1) to distinguish scRNA-seq from snRNA-seq profiles.

Gene importance was quantified using the mean decrease in impurity (MDI), calculated as the cumulative reduction in Gini impurity attributable to each gene across all decision trees. Genes were ranked according to their normalized importance scores, and the top-ranked features were subsequently evaluated using differential detection rates between modalities. Differential detection was defined as the difference between single-cell and single-nucleus detection rates. Genes were classified as single-cell– or single-nucleus–associated based on the direction of this difference, and the top 200 genes ranked by feature importance were retained as a modality-associated gene resource (Ext. Data Table 2).

### Classification of droplet modality profiles by CellOrNuc function

Droplet profiles were classified using the top 10 SC- and top 10 SN-associated genes. For each droplet, raw UMI counts were summed over the two gene sets to give *x* and *y*, and the modality fraction *p* = *x*/(*x*+*y*) was computed.

Across the whole HKCA, a two-component beta-binomial mixture (*x* | *m*, *q* ∼ Binomial(*m*, *q*); *q* | *z* = *k* ∼ Beta(*a*□, *b*□)) was fitted to *p* by expectation–maximisation algorithm, with the beta layer accommodating within-population overdispersion. Components were initialised from droplets with *p* > 0.7 and *p* < 0.3 to avoid degenerate solutions in cell types dominated by a single modality.

Droplets with posterior probability γ > 0.95 for the cell-like component were classified as cell-like and those with γ < 0.05 as nucleus-like. Profile calls were crossed with the recorded assay label to define nucleus-like cells (single-cell assay, nucleus-like profile) and cell-like nuclei (single-nucleus assay, cell-like profile, Ext. Data Fig. 7).

### Differential abundance analysis (Milo)

Differential cell type abundance across conditions was assessed using Milo^71^. A k-nearest neighbor graph was constructed from the latent embedding, and local neighborhoods (n=80) were defined to capture transcriptionally similar cells. Statistical testing for differential abundance was performed using a generalized linear model framework based on the diseases included in the HKCA, incorporating experimental sex as a covariate. P values were corrected for multiple testing using spatial false discovery rate (FDR, <0.1), and significantly enriched or depleted neighborhoods were mapped back to cell type annotations for interpretation.

### Differential gene expression and pathway enrichment analysis

Differential gene expression analysis was performed using a pseudobulk approach, aggregating counts across cells within each sample and cell type using decoupler.^120^ Statistical testing was conducted using DESeq2, with disease condition specified as the design factor. Dispersion estimation and model fitting were performed using default parameters, with Cook’s distance–based outlier detection and refitting enabled.

Differential expression statistics, including Wald test–derived t-statistics, were used to rank genes for downstream analysis. Genes were considered significant based on adjusted P value thresholds (< 0.05) and log fold-change criteria (> 1).

To interpret the functional relevance of differential expression patterns, pathway enrichment analysis was performed using decoupler and gseapy. ^121^ Gene-level statistics derived from differential expression analysis were used as input for enrichment testing against curated gene sets from MSigDB Hallmark (v2025.1). Enrichment scores were computed to identify biological processes and signaling pathways significantly associated with each condition.

### GWAS integration and single-cell disease relevance scoring

To link cell-type-specific transcriptional programs to genetic risk of kidney function, we analyzed genome-wide association study (GWAS) summary statistics for serum creatinine levels derived from a large multi-ancestry meta-analysis.^84^

Summary statistics were harmonized and processed using a custom pipeline based on the MAGMA framework.^122^ Briefly, SNPs were mapped to genes using genomic coordinates based on the GRCh37 reference genome. Linkage disequilibrium (LD) structure was estimated using the 1000 Genomes Project reference panel,^123^ which was processed using LDSC ^124^ to ensure compatibility with downstream analyses.

To resolve the distribution of genetic risk at single-cell resolution, we applied single-cell disease relevance scoring (scDRS)^86^. Gene sets derived from GWAS-associated genes were used as input, and disease relevance scores were computed for individual cells in the HKCA based on their expression profiles. Prior to scoring, a cell-level covariate matrix was constructed including an intercept term, the number of detected genes per cell, and batch indicators corresponding to individual samples and dataset of origin. These covariates were incorporated into the scDRS framework to account for differences in sequencing depth and technical variation across datasets. Disease relevance scores were computed at the single-cell level and subsequently aggregated by cell type for downstream enrichment analyses.

### Analysis of spatial transcriptomics kidney datasets using the HKCA

#### Spatial transcriptomics datasets and preprocessing

Spatial transcriptomics data were obtained from publicly available sources including 10X Genomics for the Visium HD Human kidney FFPE^52^, the non-diseased kidney of the Xenium Human Multi-Tissue panel^59^, and GSM8216268^70^. Raw spatial gene expression matrices were processed using spatialdata-io (v0.6.0) and the SpatialData package (v0.7.3a0)^125^ pipelines provided by the respective platforms. We used Cellpose^126^ to create a segmentation mask when not available with the original dataset and performed probabilistic transcript correction using Proseg.^127^ For high-resolution Visium HD data, spot-level transcriptomic profiles were further processed using the bin2cell^128^ framework to aggregate spatial barcodes into cell-like units. Both bin level data and aggregate spatial data were used for downstream analyses.

#### Cell type mapping to spatial data

Cell type identities were assigned to spatially resolved profiles using reference-based mapping from the HKCA. Gene expression signatures derived from the annotated and harmonized single-cell and single-nucleus populations were used to train a multinomial logistic regression classifier implemented in CellTypist^129^ (v.1.7.1).

For VisiumHD data, gene expression matrices at sub-spot resolution (8μm bins) and bin2cell-aggregated profiles (2μm bins) were used following quality control for cells with low transcript counts (<45 genes), normalization, and log1p transformation. The trained HKCA model was applied using the celltypist.annotate() function with probability threshold(p_thres) set at 0.5 and majority voting set to True.

For Xenium datasets, cell type annotation was performed using an optimal transport–based mapping approach as implemented in TACCO^130^, which decomposes spatial profiles into probabilistic cell type contributions, enabling transfer of atlas-derived cell type labels while accounting for gene detection differences between single-cell and targeted spatial transcriptomic data.

#### Spatial neighborhood and proximity analysis

To characterize spatial organization and cell–cell interactions, neighborhood and proximity analyses were performed using the Squidpy v(1.6.6) framework^131^. Spatial graphs were constructed based on physical proximity of cells or spatial bins, using the sq.gr.spatial_neighbors() with n_neighs=8 and neighborhood enrichment statistics were computed to identify significantly co-localizing cell type pairs by sq.gr.co_occurrence() and sq.gr.nhood_enrichment() with n_perms =1000.

Raw co-occurrence probabilities (Immune | Fibroblast) were calculated within a specified spatial radius of 50 - 1000um, followed by the computation of a spatial enrichment score defined as the log_2_ ratio of observed to expected co-occurrence frequencies. To identify preferential recruitment niches and minimize signal from non-specific interstitial background, enrichment scores were normalized across selected celltype subsets using a Z-score transformation. Network visualization was utilized to map the physical architecture of these interactions, with edge weights scaled by raw co-occurrence intensity to represent the structural connectivity of the stromal-immune bridge. Finally, the non-finite Z-score values, arising from permutation-based neighborhood enrichment analysis under conditions of low cell counts or absent interactions, were replaced with zero to represent neutral enrichment and ensure numerical stability in downstream analyses.

#### Atlas-guided annotation of the spatial kidney atlas of diabetic kidney disease and cell type abundance association with kidney function

To transfer HKCA cell type annotations to the spatial atlas,^8^ the previously trained logistic regression classifier was applied to spatial transcriptomics profiles generated using Xenium and CosMx platforms. Cell type predictions were generated using CellTypist with majority-voting enabled to improve annotation robustness and probability threshold (p_thres) set at 0.5.

Predicted labels were integrated with the original spatial atlas annotations to generate a unified cell type annotation framework. Cell type concordance between HKCA-derived predictions and author-provided annotations was assessed using contingency tables and normalized confusion matrices. Rare populations represented by fewer than 20 cells were excluded from concordance analyses. Finally, the niche previously identified by the authors were used with the transferred annotation from the HKCA for analysis.

To evaluate relationships between cellular composition and kidney function, spatial neighborhood abundances were correlated with estimated glomerular filtration rate (eGFR) categories as provided by the authors. eGFR values were grouped into ordinal categories (<30, 30–60, and >60 mL min ¹ 1.73 m ²) and encoded as ranked variables (1–3). For each disease condition and spatial niche, the relative abundance of individual cell types was correlated with eGFR rank using Spearman’s rank correlation coefficient.

Correlations were calculated independently for each cell type within a given niche and disease condition. Analyses were restricted to groups containing at least four samples and cell types exhibiting non-zero variance across samples. Spearman correlation coefficients (ρ) and associated p-values were used to quantify the strength and direction of associations between cell type abundance and kidney function.

### Trajectory and fate inference analysis

To investigate stromal state transitions, we performed trajectory inference on the fibroblast compartment using Palantir (v1.4.1)^132^. Analyses were restricted to fibroblast-lineage cells and initialized from an adventitial fibroblast root state selected on the basis of its position in diffusion space and expression of FOXD1. Diffusion maps were computed on the scVI latent embedding, and MAGIC was used to generate imputed expression values for downstream trend analyses. Palantir was run with 500 waypoints and k-nearest-neighbor parameterization (k = 50), with predefined terminal states corresponding to Norn cells, Fibroblasts_INHBA, Myofibroblasts, Fibroblasts_NEGR1□, Medullary_fibroblasts, and Perivascular_fibroblasts. Pseudotime values and branch assignments were extracted from the Palantir output and visualized on the low-dimensional embedding.

To quantify lineage commitment, transition dynamics were further modeled using CellRank.^133^ A pseudotime kernel was constructed from the Palantir pseudotime values, and a transition matrix was computed using a soft-threshold scheme. Fate probabilities were then estimated with the GPCCA estimator,^134^ enabling lineage-level assessment of terminal state probabilities across fibroblast subpopulations. Gene expression trends along the inferred trajectory were modeled using MAGIC-imputed expression values and GAMR-based smoothing, allowing identification of dynamic programs associated with lineage progression and state bifurcation.

### Ligand–receptor interaction analysis

Ligand–receptor interactions and downstream signaling effects were inferred using NicheNet.^60^ Sender and receiver cell populations were defined based on annotated cell types within the HKCA. A receiver-centered NicheNet approach was applied using all expressed genes in more than 20% cells of the receiver.

Ligand activity was prioritized based on regulatory potential scores computed by NicheNet, which integrate prior knowledge of ligand–target gene networks. Top-ranked ligands were further filtered based on expression in sender cell populations and the presence of corresponding receptors in receiver cells. Predicted ligand–target interactions were used to infer intercellular communication networks, and results were visualized to highlight key signaling pathways mediating stromal–immune and vascular interactions in both healthy and diseased contexts.

### Hormone2Cell analysis

To infer hormone responsiveness, hormone receptor expression was evaluated using the curated hormone receptor gene definitions included in the Hormone2Cell package^64^. Receptor activity scores were calculated independently for single-cell and single-nucleus data and subsequently combined to generate consensus receptor expression profiles for each annotated kidney cell type as suggested by the authors.

Potential hormone-mediated communication networks were inferred by integrating the Norn cell profiles with receptor expression across all kidney cell types and non-kidney tissues. Hormones whose receptors were expressed in distant cell populations were classified as candidate endocrine signaling interactions, whereas receptor expression in renal cell populations was considered indicative of potential paracrine communication.

### Drug-gene interaction and target enrichment analysis

To identify therapeutically relevant pathways and potential drug targets, we performed drug–gene interaction analysis using Drug2Cell.^87^ Gene expression profiles from annotated cell populations were scored against curated drug–target gene sets using CHEMBL v36 (https://www.ebi.ac.uk/chembl/), to infer drug-associated activity across cell types. Drug2Cell computes enrichment scores based on the expression of known drug target genes, enabling systematic prioritization of compounds with potential relevance to specific cellular states. These scores were used to identify drugs and small molecules associated with certain celltype populations.

#### Staining Immunohistochemistry

Kidneys donated for transplantation but subsequently deemed unsuitable were acquired through the NHS Blood and Transplant (NHSBT) service, REC12/EE/0446. Biopsies 0.5-1cm in thickness were fixed in freshly prepared 1% paraformaldehyde (ThermoFisher Scientific, cat# 28908) solution for 24 hours at 4 degrees then rinsed in PBS for 2×5 minutes before being dehydrated in a 30% sucrose solution for 24 hours at 4 degrees, embedded in OCT (CellPath, cat# KMA-0100-00A) and kept at −70 degrees until sectioning. 25μm sections were placed on Polysine slides (ThermoFisher Scientific, cat# J2800AMNZ) and left to air dry for 10 minutes at room temperature. Sections were blocked in 0.1M Tris containing 0.3% Triton-X (Sigma-Aldrich, cat# 92426-100ML), 1% BSA (R&D Systems, cat# DY995), 1% normal rat serum (ThermoFisher Scientific, cat# 10-710-C) and 1% normal donkey serum (Abcam, ab7475) for 30 minutes at room temperature. Primary antibodies were diluted in blocking solution and incubated for 1 hour at room temperature before washing in PBS for three times 10 minutes. Secondary antibodies were then prepared as above and incubated for 1 hour at room temperature before washing. When a third staining step was necessary due to conflicting antibody species (unconjugated and conjugated mouse IgG1 κ antibodies), samples were blocked again for 30 minutes with added normal mouse serum (Invitrogen, cat# 10410) and the tertiary antibodies were prepared in blocking solution containing normal mouse serum for incubation as above and washing in PBS (See full list of antibodies in Ext.Data table 2). Slides were then mounted in 20μL Fluoromount-G (SouthernBiotech, cat# 0100-01). Images were acquired on a Leica SP8 confocal microscope using a 40x 1.1N/A water immersion objective and later processed on Imaris (Bitplane). Information for all antibodies used can be found in Ext. DataTable 6.

### Urinary and mice single cell prediction from the HKCA

Using scArches-based reference mapping^97^, we projected urine-derived or murine derived single-cell transcriptomes onto the HKCA, enabling consistent annotation of epithelial, stromal, and immune populations across independent cohorts. The pretrained scVI reference model was used as the foundation for transfer learning, enabling projection of query datasets into the HKCA latent space while preserving the reference structure. Query models were trained for up to 400 epochs using a batch size of 256, mixed-precision GPU training, and a learning rate of 5 × 10^-4^. Model optimization employed early stopping based on validation ELBO (patience = 25 epochs).

The urine datasets were downloaded from GEO (GSE176465, GSE193512, GSE199321, GSE282344), while the murine dataset was downloaded from the IGVF data portal (https://data.igvf.org/) and the ischaemia reperfusion murine datasets were downloaded from GEO (GSE180420, GSE182256, GSE139107, GSE190887). Following initial quality control for low counts, mitochondrial and ribosomal genes mapping, cell type identities were transferred from the HKCA reference trained model with scArches to query cells using nearest-neighbor relationships within the shared latent space. Prediction confidence scores were used to assess annotation reliability, and mapped annotations were used for downstream compositional and disease-stratification analyses. Concordance between transferred labels and author-provided annotations, was used to evaluate annotation performance, while the mapped gene expression space was inspected to identify cell populations not captured in the original annotations.

Dimensionality reduction of log-transformed cell type proportions by principal component analysis (PCA) demonstrated stratification of patients in the urine cohort into separable groups, broadly distinguishing glomerular from inflammatory disease states. These axes of variation were driven by coordinated changes in immune activation and tubular epithelial composition, consistent with underlying disease biology. Statistical comparisons of cell type abundances across murine strains were conducted using the Kruskal-Wallis test, followed by Dunn’s post-hoc test with Benjamini-Hochberg FDR correction for multiple comparisons.

## Code availability

Code used for the integration and single cell atlas analysis can be found on https://github.com/Peng-He-Lab/Kidney_integration. The trained models from Celltypist and scArches on the HKCA are available at Zenodo (https://zenodo.org/records/21468481).

## Data availability

The diabetic kidney disease spatial datasets can be downloaded at https://zenodo.org/records/15007208. The raw and processed data for the mouse strain study are publicly available through the IGVF Data Portal (https://data.igvf.org/). The murine ischaemia reperfusion datasets can be downloaded from GEO (<u>GSE180420</u>, <u>GSE182256</u>, <u>GSE139107</u>, <u>GSE190887</u>). The urine datasets are available at NCBI (<u>GSE176465</u>, <u>GSE199321</u>, <u>GSE199321</u>, <u>GSE282344</u>). The HKCA Integrated objects are available at the CellxGene data portal and Cell Annotation Platform (https://doi.org/10.57772%2Fcap.lr1qubc8lvq53byicflpmnedecs2.1006).

## Acknowledgements

This work was supported by the Chan Zuckerberg Initiative via grant CZIF2022-007488 - HCA Data Ecosystem. S.A.T., M.D.L is supported by the CZI data ecosystem grant. P.H. is supported by the Esther Simon Memorial Fund from the Research Evaluation and Allocation Committee at UCSF, and Sandler Program for Breakthrough Biomedical Research, which is partially funded by the Sandler Foundation. K.S. is supported by the NIHR Academic Clinical Fellowship ACF-2021-14-013. P.H, K.S and W.Z are supported by UC Noyce Initiative Award. The Kidney Precision Medicine Project (KPMP) is supported by the National Institute of Diabetes and Digestive and Kidney Diseases (NIDDK) through the following grants: U01DK133081, U01DK133091, U01DK133092, U01DK133093, U01DK133095, U01DK133097, U01DK114866, U01DK114908, U01DK133090, U01DK133113, U01DK133766, U01DK133768, U01DK114907, U01DK114920, U01DK114923, U01DK114933, U24DK114886, UH3DK114926, UH3DK114861, UH3DK114915, UH3DK114937, 1U01DK144965-01, 1U01DK144994-01. We gratefully acknowledge the essential contributions of our patient participants and the support of the American public through their tax dollars.” The authors acknowledge the University of Michigan Medical School Central Biorepository (RRID:SCR_026845) for providing biospecimen storage, management, and distribution services in support of the research reported in this publication/grant application/presentation. The content is solely the responsibility of the authors and does not necessarily represent the official views of the National Institutes of Health. This publication is part of the Human Cell Atlas.

## Author contributions

KS, HW, AP, RT, NR, MT designed and performed the experiments, contributed to the analysis and experimental design. KS and PH wrote the manuscript. All authors reviewed the manuscript.

## The Human Cell Atlas Kidney Bionetwork

Elnaz Abollahzadeh18, Andrea Califano19,20,21 Sarah Q. Crome22,23,24,25 Christian Krebs26, Blue Lake27, Siyu Lin,28,29,30,31 Ali Mortazavi,18 Julia M. Murphy,24,25 Aleksandar Obradovic32, Ben Stewart5, 6, Poorvita Vijayananda1, Aijun Wang28,29,30,31 Kun Zhang27,33, Yuping Zhang34, Yu Zhao35

18. Department of Developmental and Cell Biology, University of California, Irvine, Irvine, CA, USA
19. Department of Systems Biology, Columbia University Irving Medical School, New York, NY, United States of America
20. Chan Zuckerberg Biohub New York, New York, NY, USA;
21. Departments of Biochemistry & Molecular Biophysics, Medicine, and Biomedical Informatics, Columbia University Irving Medical Center, New York, NY, USA
22. Terry Fox Laboratory, Basic and Translational Research, BC Cancer Research Institute, Canada
23. University of British Columbia, Faculty of Medicine, Department of Medical Genetics, Canada
24. University Health Network, Toronto General Hospital Research Institute, Ajmera Transplant Centre, Canada
25. University of Toronto, Temerty Faculty of Medicine, Department of Immunology, Canada
26. Hamburg Center for Translational Immunology (HCTI), University Medical Center Hamburg-Eppendorf, Hamburg, 20246, Germany
27. San Diego Institute of Science, Altos Labs, San Diego, CA, USA
28. Center for Bioengineering in Medicine, School of Medicine, University of California Davis, Sacramento, CA
29. Department of Surgery, School of Medicine, University of California, Davis, Sacramento, CA
30. Department of Biomedical Engineering, University of California Davis, Davis, CA
31. Shriners Children’s Northern California, Sacramento, CA
32. Department of Medicine, Columbia University Irving Medical Center, New York, NY, USA
33. Department of Bioengineering, University of California, San Diego, La Jolla, CA, USA;
34. Institute of Medical Systems Biology, University Medical Center Hamburg-Eppendorf, Hamburg, Germany
35. Michigan Center for Translational Pathology, University of Michigan, Ann Arbor, MI, USA

## Declarations

S.A.T. is a scientific advisory board member of Bioptimus, ForeSite Labs, Xaira Therapeutics, a co-founder, Board observer and equity holder of TransitionBio, a co-founder, consultant and Board Director of Ensocell Therapeutics, a non-executive director of 10x Genomics and a part-time employee of GlaxoSmithKline. J.C.M. has been an employee of Genentech since September 2022. M.D.L. contracted for the Chan Zuckerberg Initiative and consults for CatalYm GmbH. M.K. reports grants and contracts through the University of Michigan outside of this work from AstraZeneca, NovoNordisk, Eli Lilly, Boehringer-Ingelheim, European Union Innovative Medicine Initiative, Certa Therapeutics, RenalytixAI, Regeneron

## Extended Figures

**Extended Figure 1.**
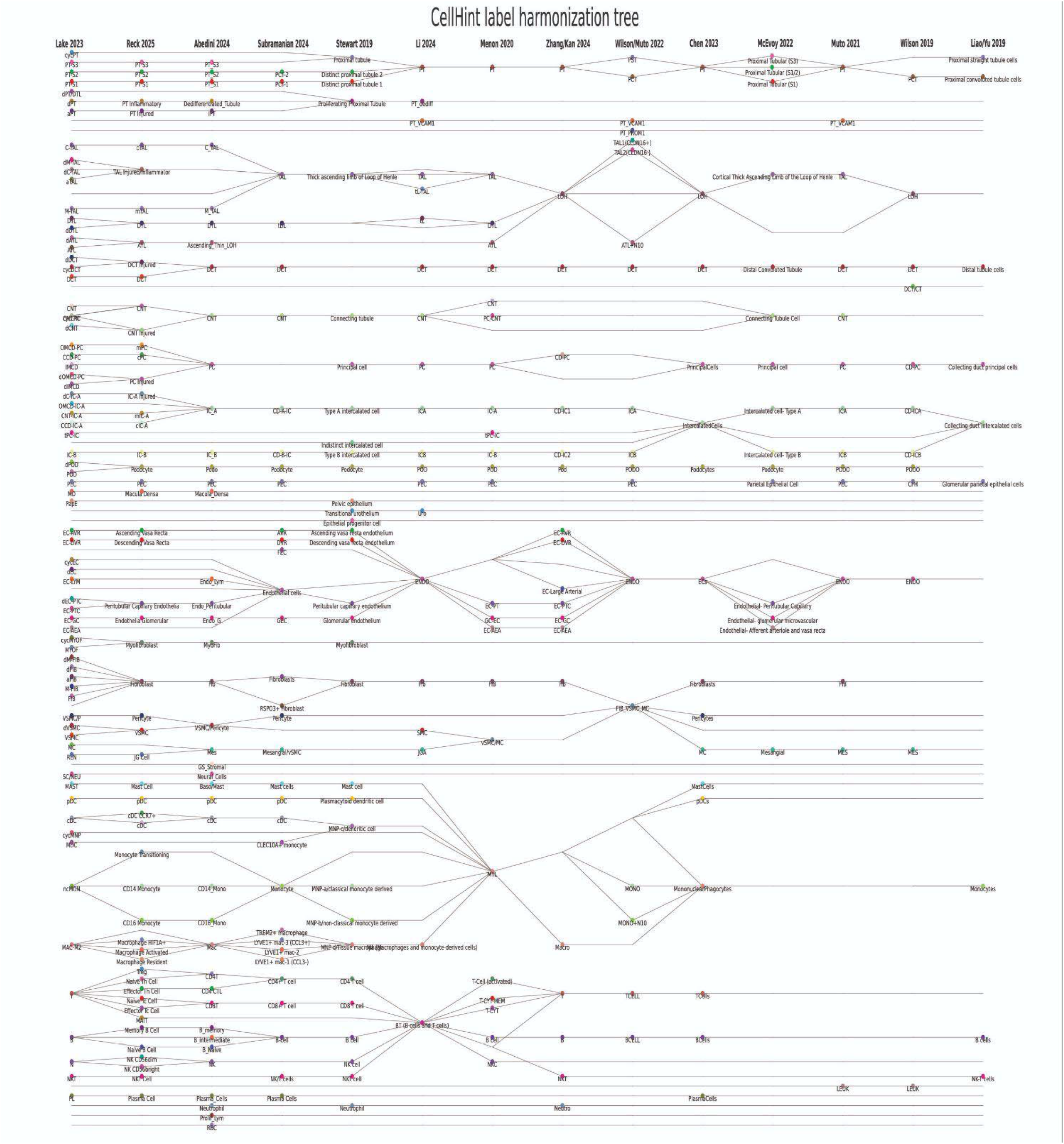
Harmonization of cell types amongst published kidney single cell datasets. Harmonization of cell type annotations amongst kidney single cell datasets was performed using CellHint^119^. The CellHint tree suggests lack of harmonized nomenclature in cell type annotation across datasets.

**Extended figure 2.**
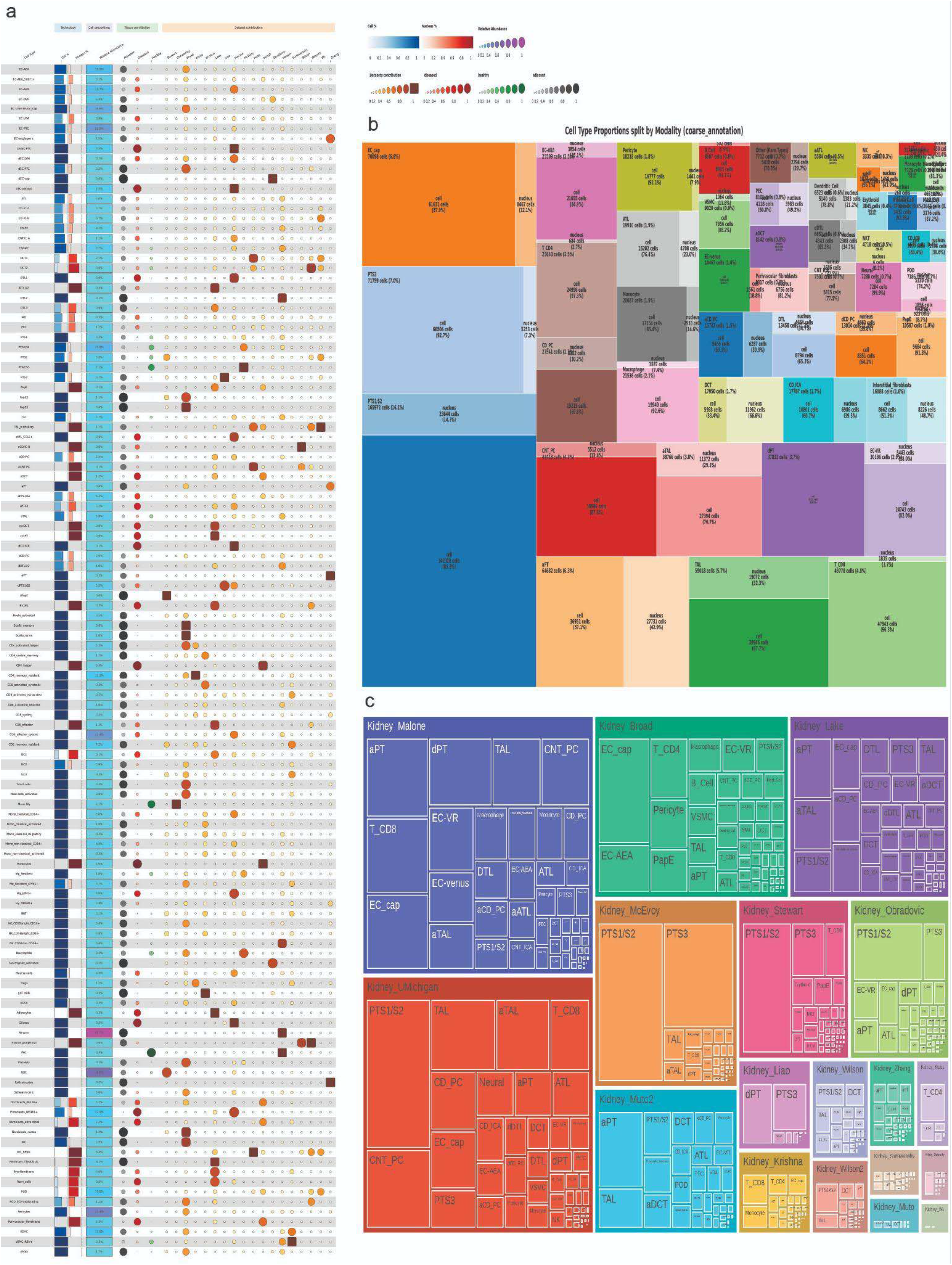
Contribution of modality and dataset per fine annotated cell type. a) Relative abundance is calculated per broader compartment epithelial, immune, endothelial stromal and other. The tissue contribution is based on the sampling origin either healthy, diseased or adjacent to tumor tissue. b) Treemap plot depicting proportion of cell types identified in the HKCA (The size of the rectangles is proportional to the size of the cell type in the atlas and contribution of cell or nucleus) c) Treemap plot depicting proportion of cells contributed by the 18 datasets comprising the HKCA (The size of the rectangles is proportional to the cell contribution of the dataset)

**Extended Figure 3.**
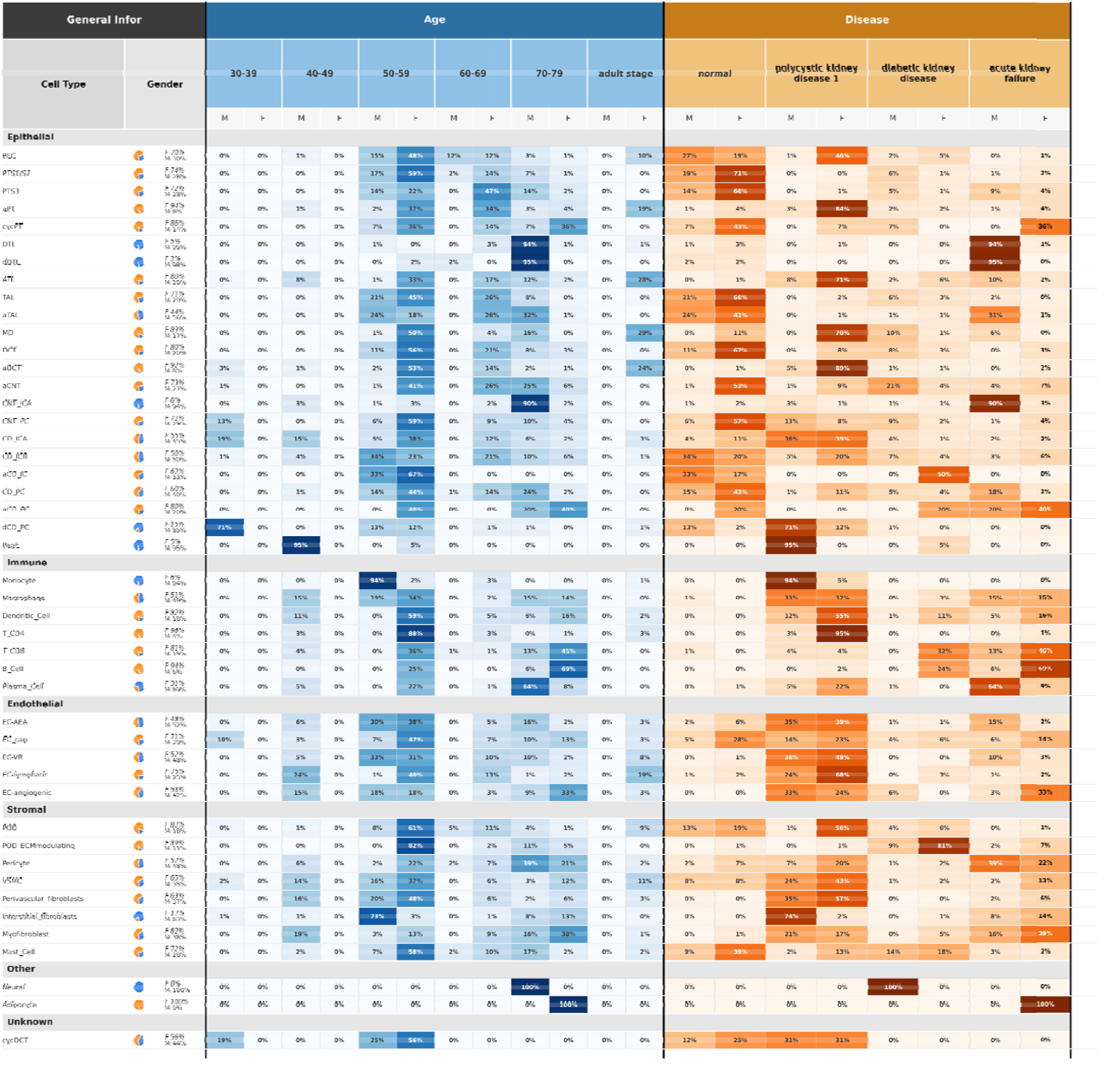
Contribution per cell type for age, gender and diseases included in HKCA single nucleus. Gender contribution per cell type denoted by the blue (male - M) and orange (female - F) in the pie chart. Age contribution per cell type and sex (if not available broader adult stage label denotes age above 20 years of age). Disease contribution per cell type and sex for each of the three diseases included in the single cell HKCA acute kidney failure, polycystic kidney disease and diabetic kidney disease.

**Extended Figure 4.**
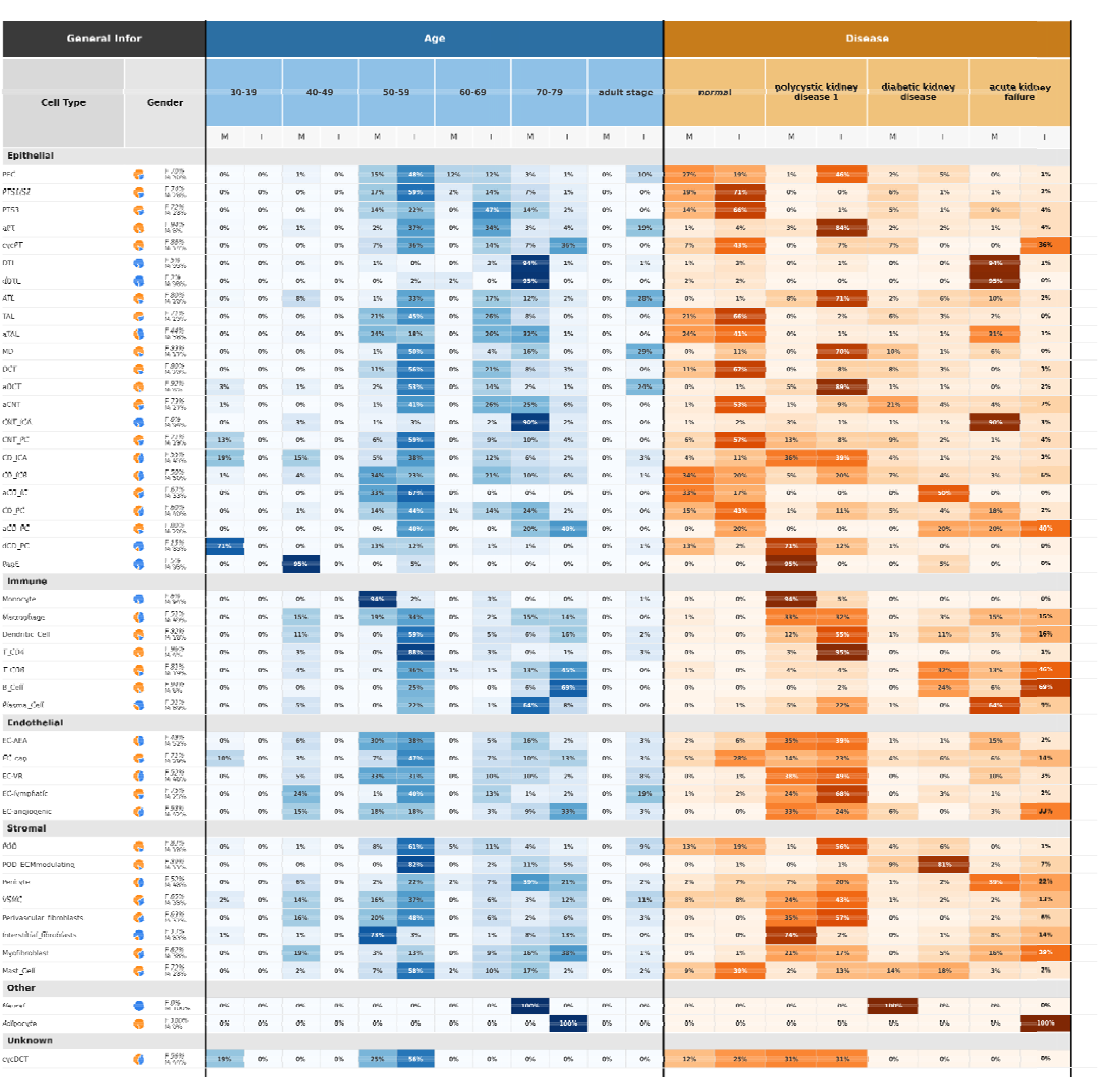
Contribution per cell type for age, gender and diseases included in HKCA single cell. Gender contribution per cell type denoted by the blue (male - M) and orange (female - F) in the pie chart. Age contribution per cell type and sex if not available broader adult stage label denotes age above 20 years of age. Disease contribution per cell type and sex for each of the three diseases included in the single cell HKCA acute kidney failure, ANCA positive glomerulonephritis and transplant rejection.

**Extended figure 5.**
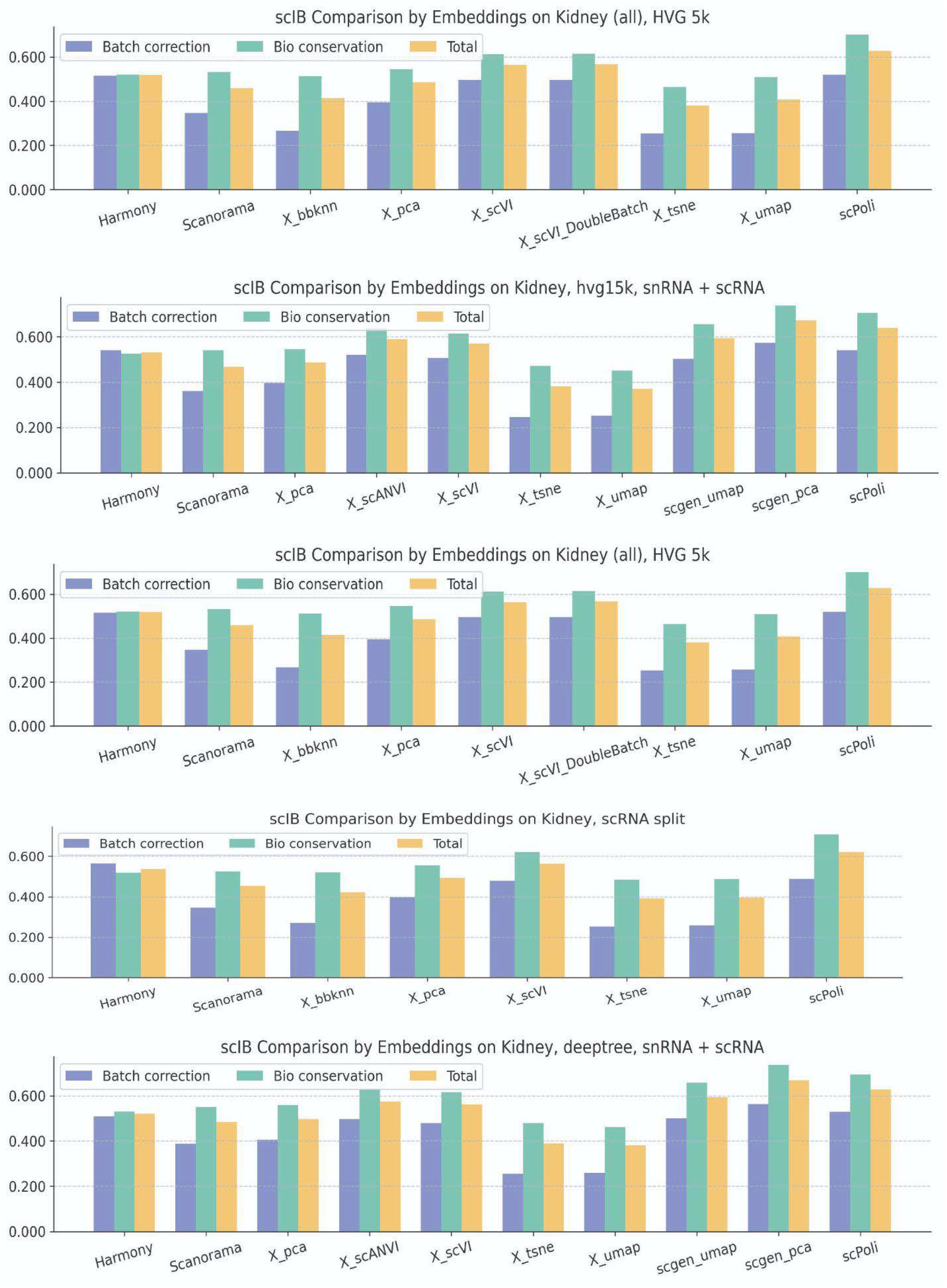
scIB metrics for different combinations of preprocessing and integration methods. Integration metrics have been calculated based on different feature selection methods high variable genes (HVG) as implemented in scanpy, or Deeptree algorithm for denoising. Similarly, cutoff for 5000 or 15000 HVG have been used to evaluate integration and also metrics have been calculated for both modalities (sc and sn) together or separately. The core dataset was independently annotated and therefore label aware integration based on scANVI, scPoli, and scGen was evaluated along the label unaware methods. Label aware methods are able to increase over integration methods nevertheless shows that harmonization biases may be contributed to improved performances.

**Extended figure 6.**
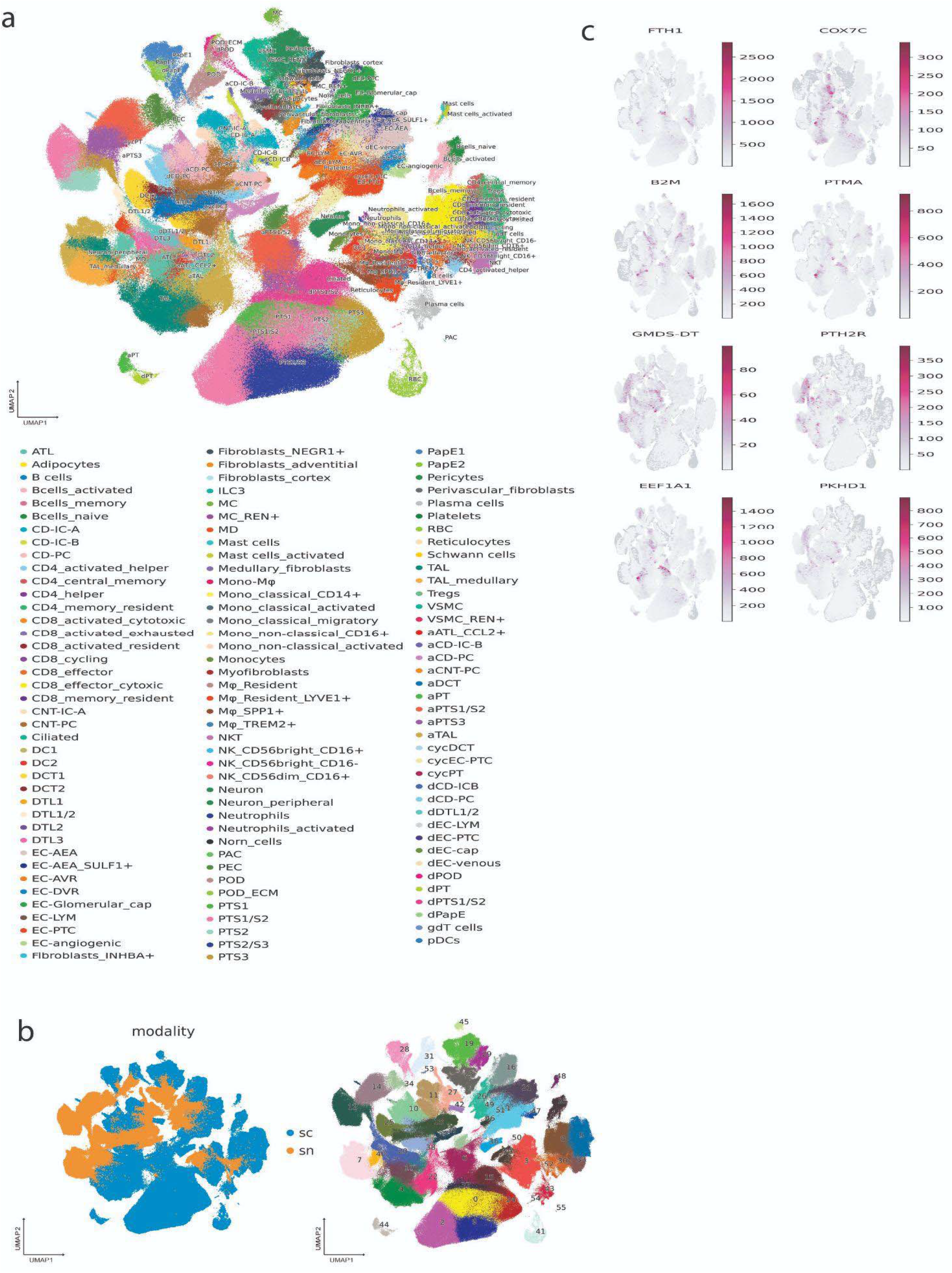
Systematic integration and technical benchmarking of single-cell and single-nucleus kidney transcriptomes. a) UMAP embedding of the integrated multi-modal dataset, annotated by fine-resolution cell types across the diverse epithelial, endothelial, myeloid, and lymphoid compartments. b) UMAP projections evaluating dataset alignment. *Left:* Cells colored by suspension type (single-cell vs. single-nucleus), demonstrating partial mixing across the latent space. *Right:* Unsupervised Leiden clustering of the integrated manifold. c) Feature plots depicting the log-normalized expression of the top technology-specific genes. Cytosolic and translational markers (e.g., *FTH1*, *COX7C*, *EEF1A1*) are heavily biased toward whole-cell capture, while long non-coding RNAs and genes with large intronic footprints (e.g., *GMDS-DT*, *PKHD1*) are enriched in single-nucleus.

**Extended Figure 7.**
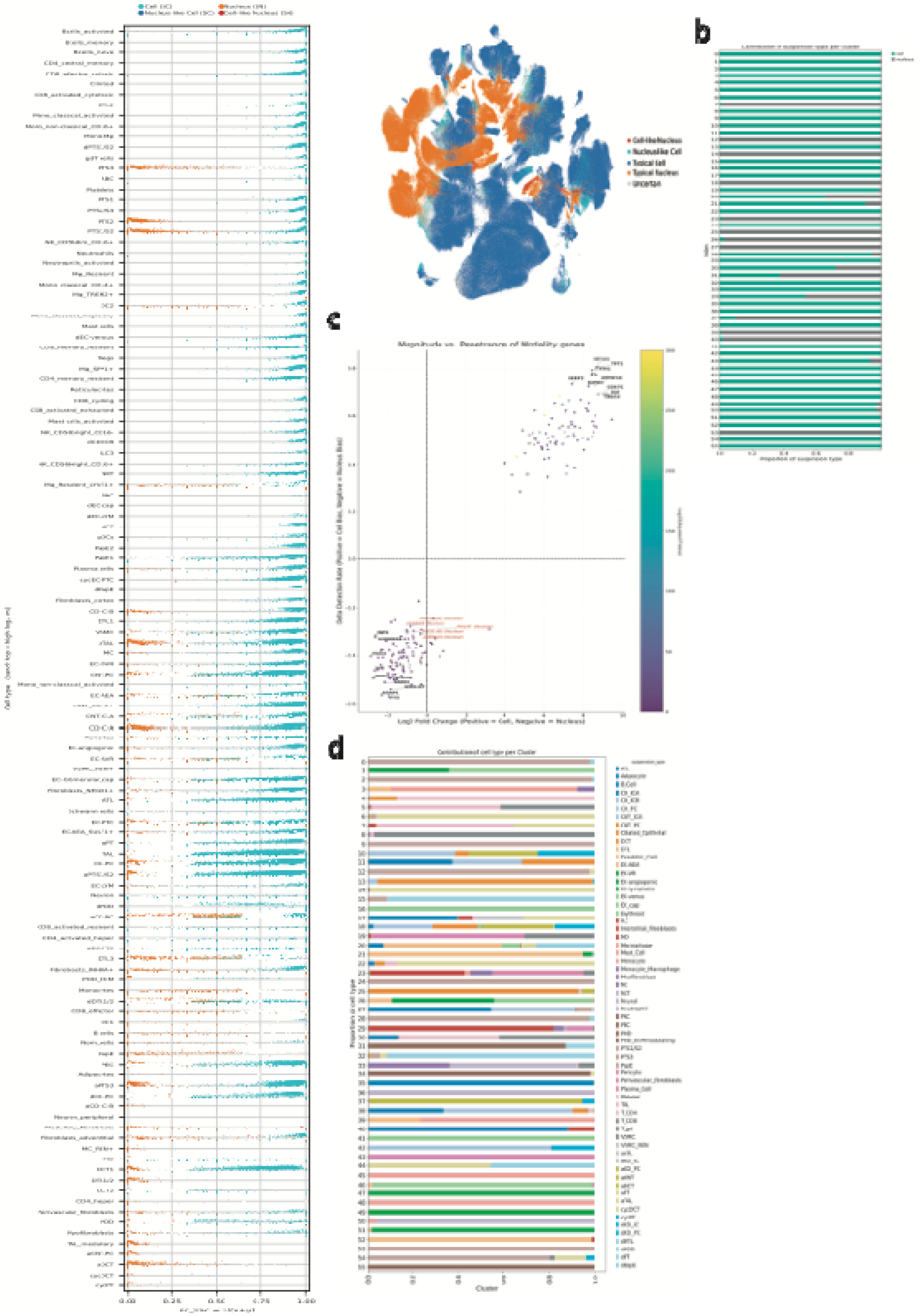
Transcriptional scoring of single-cell and single-nucleus data identifies a crossover between nucleus-like and cell-like states. a) Distribution of modality-specific signature scores in the concatenated single-cell (HKCA-cell) and single-nucleus (HKCA-nucleus) atlas. Modality assignment across the atlas. For each droplet, raw UMI counts were summed over the top 10 SC-enriched genes (*x*) and the top 10 SN-enriched genes (*y*), and the modality fraction *p* = *x*/(*x*+*y*) was computed; Across the atlas, a two-component beta-binomial mixture was fitted to *p* by expectation–maximisation algorithm, predicting a cell-like and a nucleus-like profile. Each droplet receives a posterior probability of belonging to the cell-like component, and confidently assigned profiles were crossed with the known assay label to identify cross-modality profiles (nucleus-like cell and cell-like nucleus). b) Stacked bar chart detailing the proportional contribution of single-cell and single-nucleus capture within each unsupervised Leiden cluster. c) Magnitude versus penetrance reveals distinct modality signatures in matched kidney scRNA-seq and snRNA-seq. Scatter plot comparing differential abundance (PyDESeq2 shrunken Log2 Fold Change; x-axis) and differential detection frequency as identified by the random forest classifier (Δ Detection Rate; y-axis). Positive values indicate scRNA-seq enrichment; negative values indicate snRNA-seq enrichment. Point color denotes statistical significance from the differential abundance (Log10 adjusted p-value). Annotated sub-populations highlight specific technical and biological biases: (Black/bold) Strong technical markers driven by cytoplasmic retention in whole cells (e.g., *EEF1A1*) or intronic/nuclear retention in isolated nuclei (e.g., *PKHD1*). (Red/italic) "Nuclear Bias" transcripts, primarily non-coding and antisense RNAs with high detection bias relative to overall abundance shift (|Log2FC| < 1.0; |Δ Detection| > 0.25). d) Stacked horizontal bar chart illustrating the proportional breakdown of established cell-type annotations within each Leiden cluster. We observed that ‘bridge’ clusters—such as Cluster 7, which contains a transcriptomic blend of TAL and CNT cells—exhibited substantial contributions from both single-cell and single-nucleus datasets. This indicates that complex transcriptomic states driven by physical cell-cell adhesion (in whole-cell suspensions) or ambient RNA contamination (in nuclear isolations) yield highly reproducible composite signatures. The successful co-clustering of these multi-lineage profiles demonstrates that the integration algorithm maps unique underlying transcriptional states irrespective of their biological or artifactual origin, while the incomplete mixing observed in the proximal tubular compartment highlights the limits of manifold alignment when faced with profound, capture-induced transcriptomic divergence.

**Extended Figure 8.**
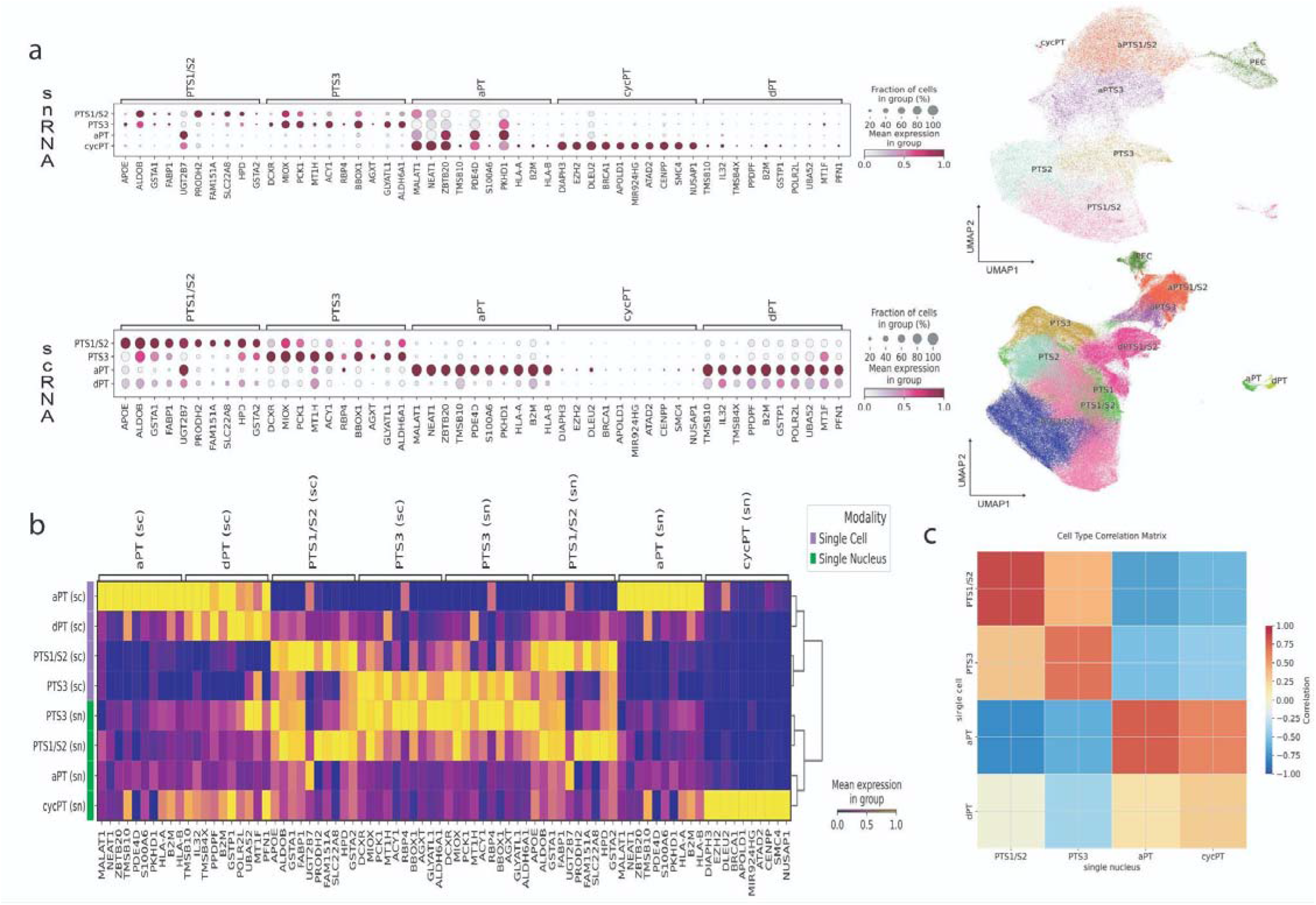
Cross-modality concordance of renal cell proximal tubule type signatures. a) Dot plots illustrating the expression of shared marker genes across single-cell (sc) and single-nucleus (sn) modalities with associated umaps for reference. Top panel displays mean expression and percentage of cells expressing marker genes in the single-nucleus dataset, while the bottom panel represents the same gene set in the single-cell dataset. Genes were selected based on statistical significance (padj <0.05) and high expression levels common to both modalities (log_2_FC >1). Cell types shown include Proximal Tubule Segments (PTS1/S2, PTS3), alteredPT (aPT), cycling PT (cycPT), and damaged PT (dPT). b) Scaled heatmap of common gene signatures by modality. Heatmap showing the mean expression of top markers scaled independently within each modality (sn, green; sc, purple) to account for differences in dynamic range between nuclear and cytoplasmic transcripts. Columns represent specific marker genes grouped by their target cell type, demonstrating high specificity and conserved expression patterns across both sequencing technologies. c) Heatmap depicting the Pearson correlation coefficients based on the expression of the shared signature gene set. High positive correlation (red) along the diagonal confirms that clusters with harmonized names (e.g., PTS1/S2 in sn vs. sc) represent the same biological populations across both datasets, validating the transfer of cell type identities.

**Extended Figure 9.**
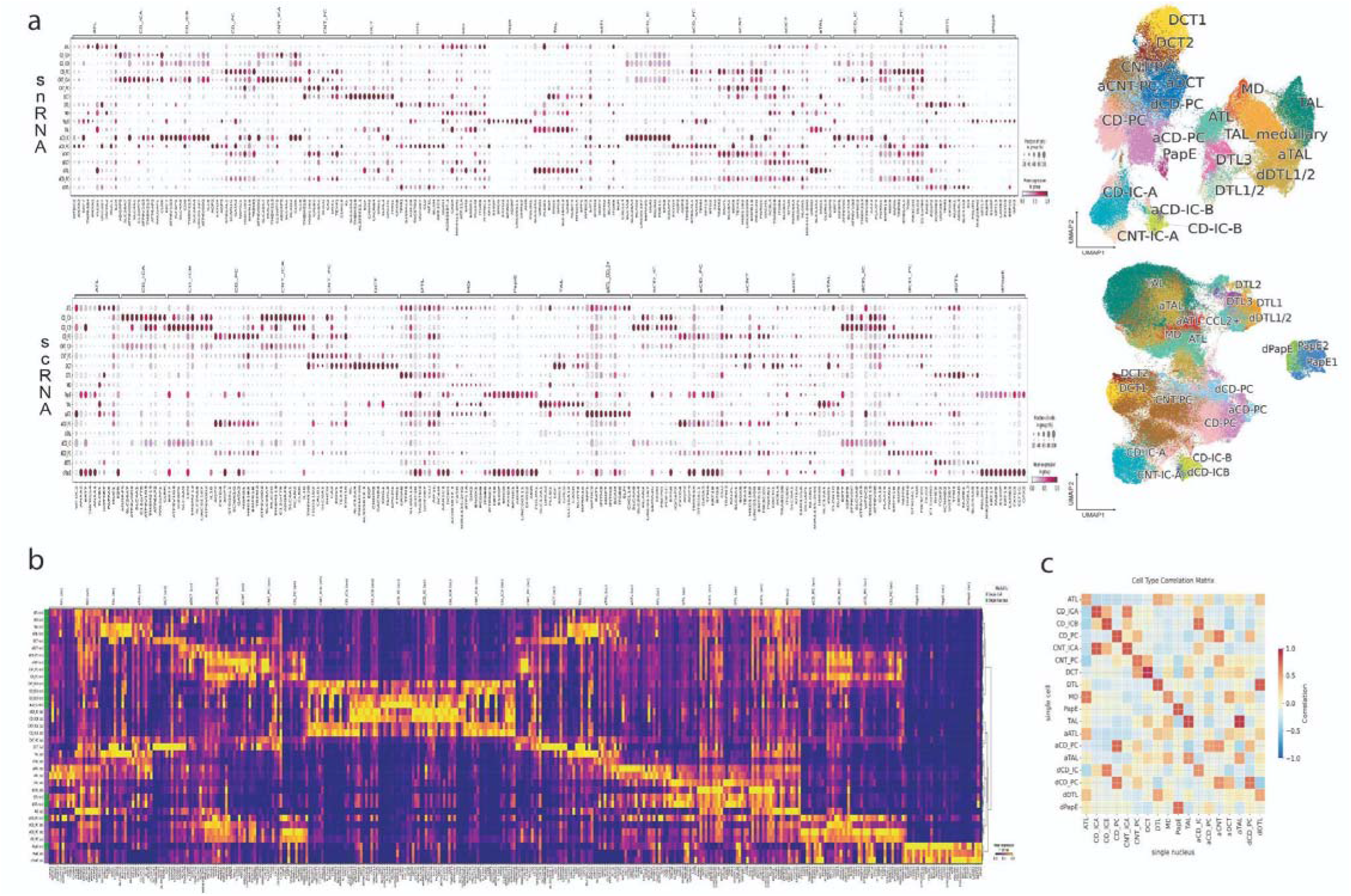
Cross-modality concordance of distal nephron and collecting duct lineages. a) Dot plots of conserved marker gene expression for distal and collecting duct cell types with associated UMAPs for reference. The top panel (sn) and bottom panel (sc) display the percentage of cells and mean expression levels for canonical and de novo markers. Distal segments: Robust expression of *SLC12A3* identifies the Distal Convoluted Tubule (DCT), while *SLC12A1* (NKCC2) and UMOD define the Thick Ascending Limb (TAL) populations. Collecting Duct (CD) System: Principal cells (CD_PC) are marked by *AQP2* and *FXYD4*, while Intercalated cells (CD_IC) show specific enrichment of *ATP6V1B1* and *CA2* (Type A, CD_ICA) or *SLC26A4* (Type B, CD_ICB). b) Integrated cross-modality heatmap of distal signature genes. Gene expression is scaled by modality (sn, green; sc, purple) to highlight the preservation of the transcriptional "fingerprint" across platforms. The heatmap demonstrates clear modularity, with distinct gene blocks corresponding to the Loop of Henle, DCT, and the various specialized cells of the collecting duct (PC vs. ICA vs. ICB). c) Correlation matrix of distal and collecting duct identities. Pearson correlation analysis between the scRNA-seq and snRNA-seq datasets. The strong diagonal confirms high concordance for all major distal cell types. Notably, the adaptive and damaged states show moderate cross-correlation, reflecting their shared transcriptional response.

**Extended Figure 10.**
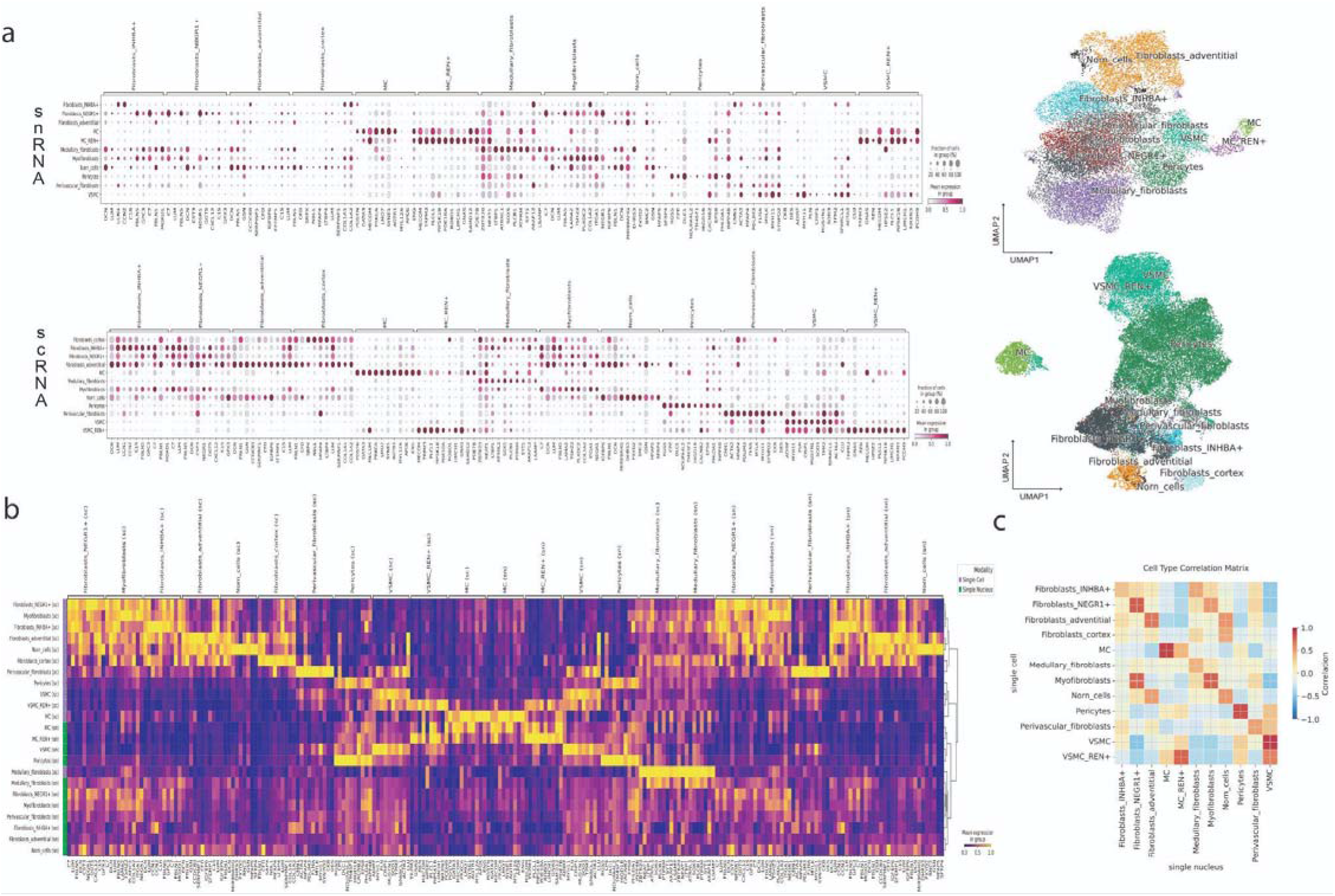
Cross-modality concordance of stromal lineages. a) Dot plots highlighting conserved marker expression in stromal sub-populations and associated UMAPs for reference. Fibroblast Specialization: Fibroblasts_INHBA^+^: Defined by an "immediate-early" signature including *CCN1*, *CCN2*, and *SERPINE1*. Fibroblasts_NEGR1^+^: Characterized by *NEGR1*, *FBLN5*, and the homeobox factor *TSHZ2*. The expression of *CXCL12* and *PDGFRA* identifies these as a key signaling hub in the interstitium. Adventitial and cortical Fibroblasts: Fibroblasts_adventitial are distinguished by PI16-related markers (CCDC80, FBLN1). In contrast, fibroblasts_cortex express a high-volume basement membrane signature (*COL1A1*, *COL1A2*, *LUM*, *MFAP4*). Endocrine and specialized interstitial Cells: Norn cells: These cells show a distinct "protective" and growth-factor-rich profile, marked by *GSN* (Gelsolin), and *CXCL14*. The high expression of *DCN* (Decorin) and *APOD* suggests a specialized niche in the peritubular microenvironment. Renin-Lineage Cells: Two distinct Renin-associated populations are resolved. MC_REN^+^ (Mesangial-like) and VSMC_REN^+^ (Smooth muscle-like) both express REN and the transcription factor *MECOM* but are distinguished by their primary lineage markers (e.g., *MYH11* in the VSMC-derived group) along with the lack of *PIEZO2* expression in the VSMC (see Figure 2). b) Scaled heatmap of stromal signature genes across modalities. Gene expression is scaled within each modality (sn, green; sc, purple). The heatmap demonstrates that the Medullary_fibroblasts possess a highly divergent signature (e.g., *ZBTB20, SOX5, HPSE2*) compared to cortical subsets, likely reflecting the extreme osmotic and hypoxic conditions of the renal medulla. c) Correlation matrix of stromal identities confirms high concordance for all major clusters. Notably, the Myofibroblasts show high similarity to the NEGR1^+^ population, while MC (Mesangial Cells) maintain a unique identity defined by GATA3 and POSTN, separate from other interstitial fibroblasts and VSMC.

**Extended Figure 11.**
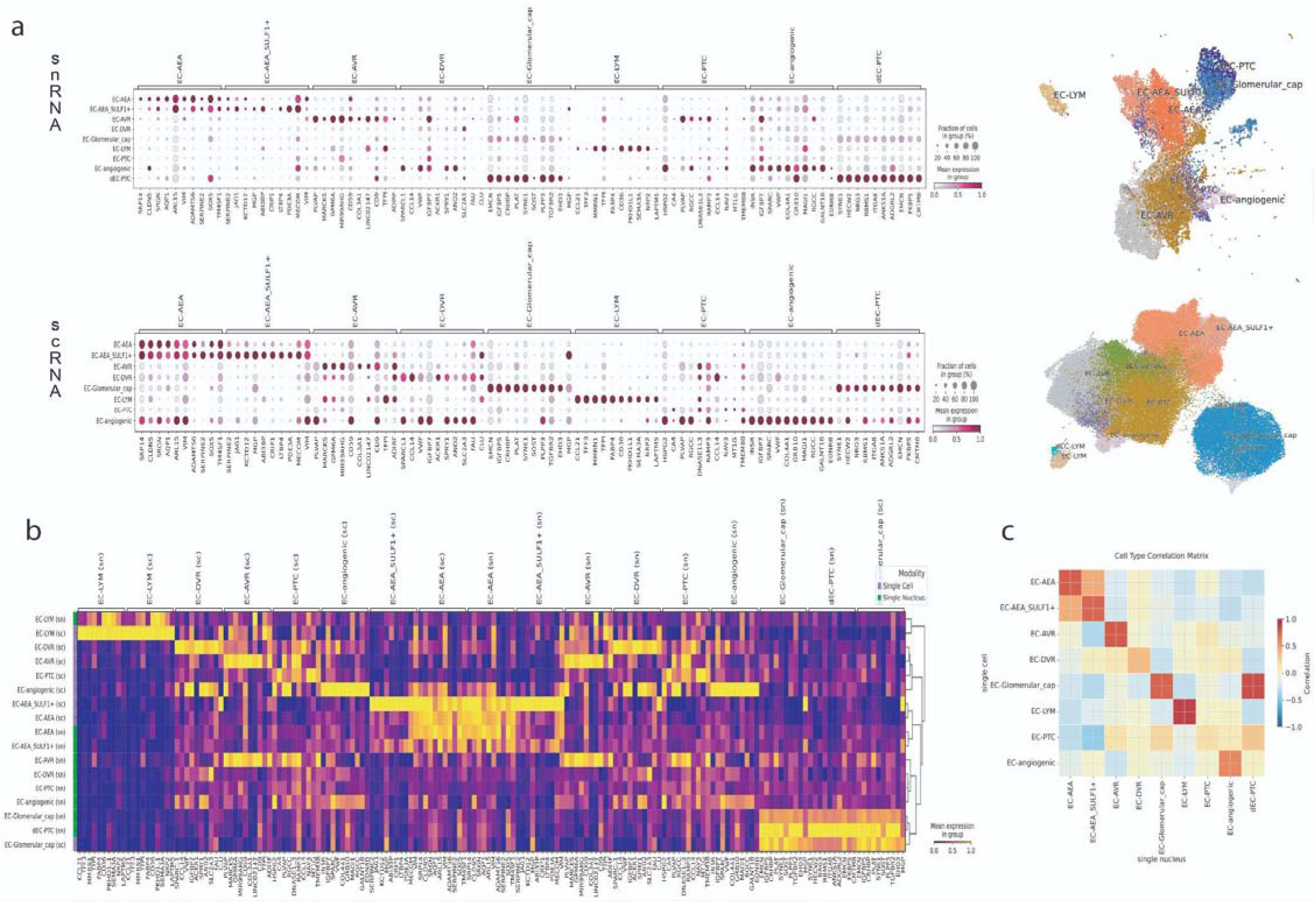
Cross-modality concordance of endothelial lineages. a) Dot plots of conserved marker gene expression across endothelial sub-lineages with associated UMAPs for reference. High-Resistance Vessels: EC-AEA (Afferent/Efferent Arterioles) are defined by arterial markers such as JAG1 and SERPINE2. A specific subset (EC-AEA_SULF1+) expresses *SULF1*, a marker associated with mechanical shear stress and specialized heparan sulfate remodeling at the pre-glomerular interface. Fenestrated Capillaries: EC-PTC (Peritubular) and EC-AVR (Ascending Vasa Recta) populations are characterized by high levels of *PLVAP*, reflecting the high-permeability requirements of adult renal reabsorption. Medullary Specialization: The EC-DVR (Descending Vasa Recta) is uniquely identified by the *GLUT3* transporter *SLC2A3*, vital for the medullary concentrating mechanism. b) Integrated cross-modality heatmap of endothelial signature genes. Gene expression is scaled by modality (sn, green; sc, purple) to demonstrate the consistency of sub-type-specific gene blocks. The high overlap in gene enrichment between modalities validates the categorization of specialized niches, such as the SULF1+ arteriole subset and the developing peritubular network. c) Correlation matrix of endothelial identities. High correlation on the diagonal confirms that the specialized identities of the renal vasculature (e.g., glomerular capillaries vs. vasa recta) are highly conserved between single-cell and single-nucleus modalities, despite differences in transcript recovery.

**Extended Figure 12.**
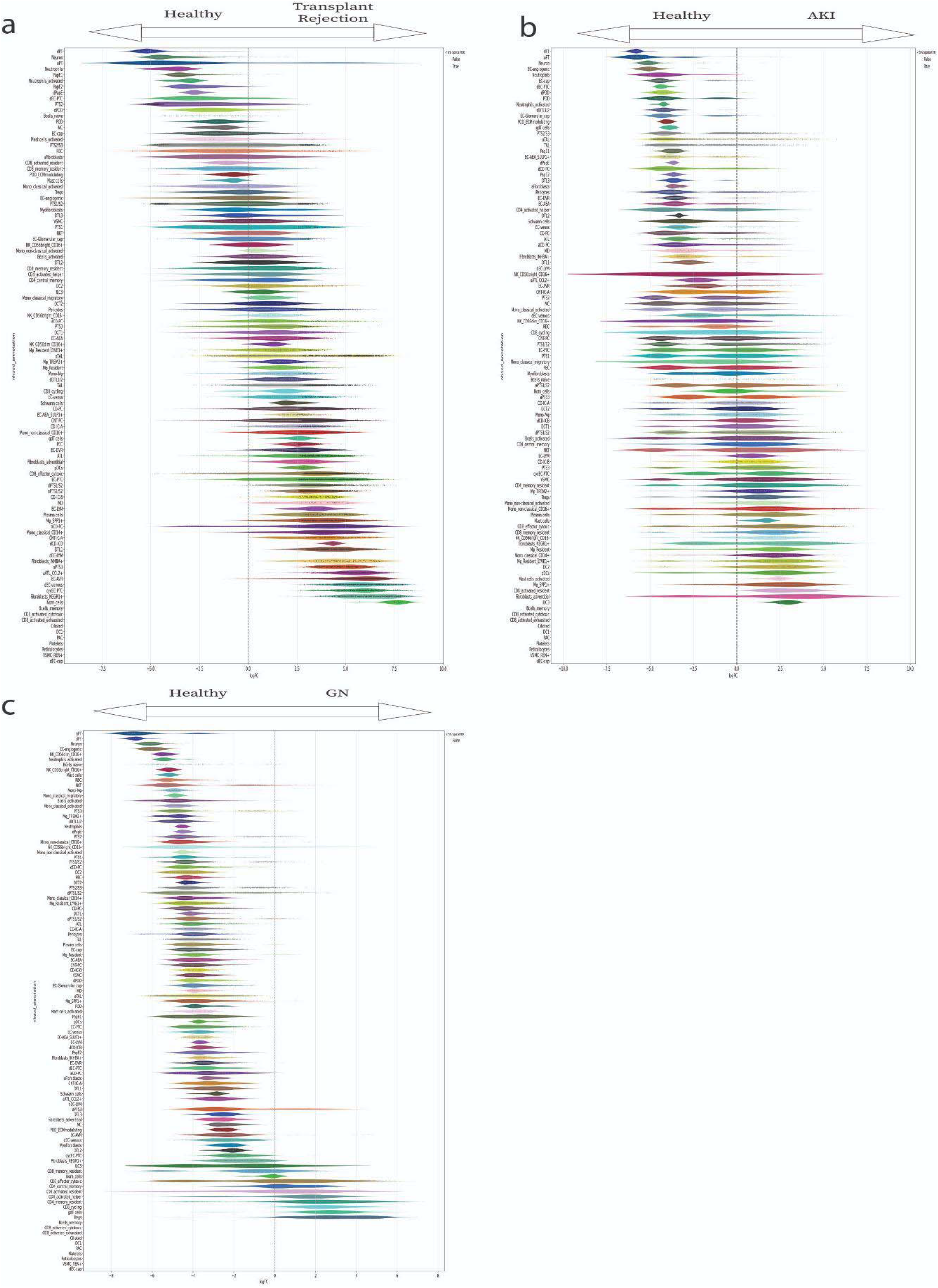
Milo analysis for cell abundance for each disease compared to healthy kidney included in the HKCA single cell. a) Cellular abundance difference between rejecting kidney transplants and normal kidney showing increased neighborhoods (spatial FDR<0.1 – black dots) in immune cells along with damage responsive aATL-CCL2^+^ and kidney immune attracting fibroblasts NEGR1^+^. b) Cellular abundance difference between acute kidney failure (AKI) and normal kidney showing increased neighborhoods (spatial FDR>0.1) in SPP1+ macrophages and minimal changes in epithelial compartments. c) Cellular abundance difference between ANCA^+^ glomerulonephritis and normal kidney showing increased neighborhoods in adaptive immune cells CD4 and CD8 T cells but with no compartment reaching statistical significance.

**Extended Figure 13.**
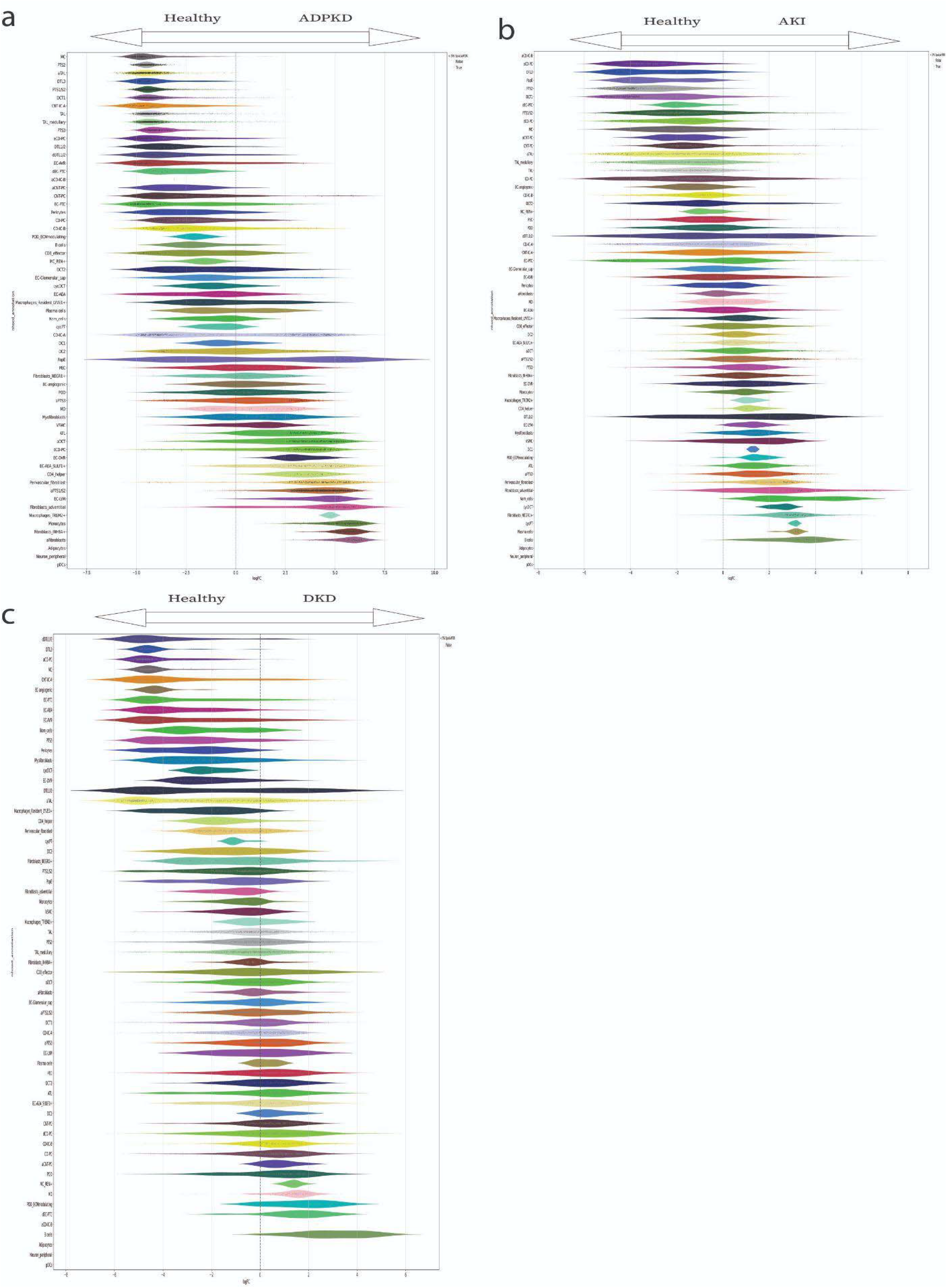
Milo analysis for cell abundance for each disease compared to healthy kidney included in the HKCA single nucleus. a) Cellular abundance difference between adult polycystic kidney disease (ADPKD) and normal kidney showing increased neighborhoods (spatial FDR >0.1 – black dots) in stromal cells along with damage responsive aPT proximal tubular cells. b) Cellular abundance difference between acute kidney failure (AKI) and normal kidney showing increased neighborhoods (spatial FDR>0.1 in epithelial compartments. c) Cellular abundance difference between diabetic kidney disease (DKD) and normal kidney showing increased neighborhoods of damaged endothelial cells in peritubular capillaries and POD_ECM podocytes but with no compartment reaching statistical significance.

**Extended Figure 14.**
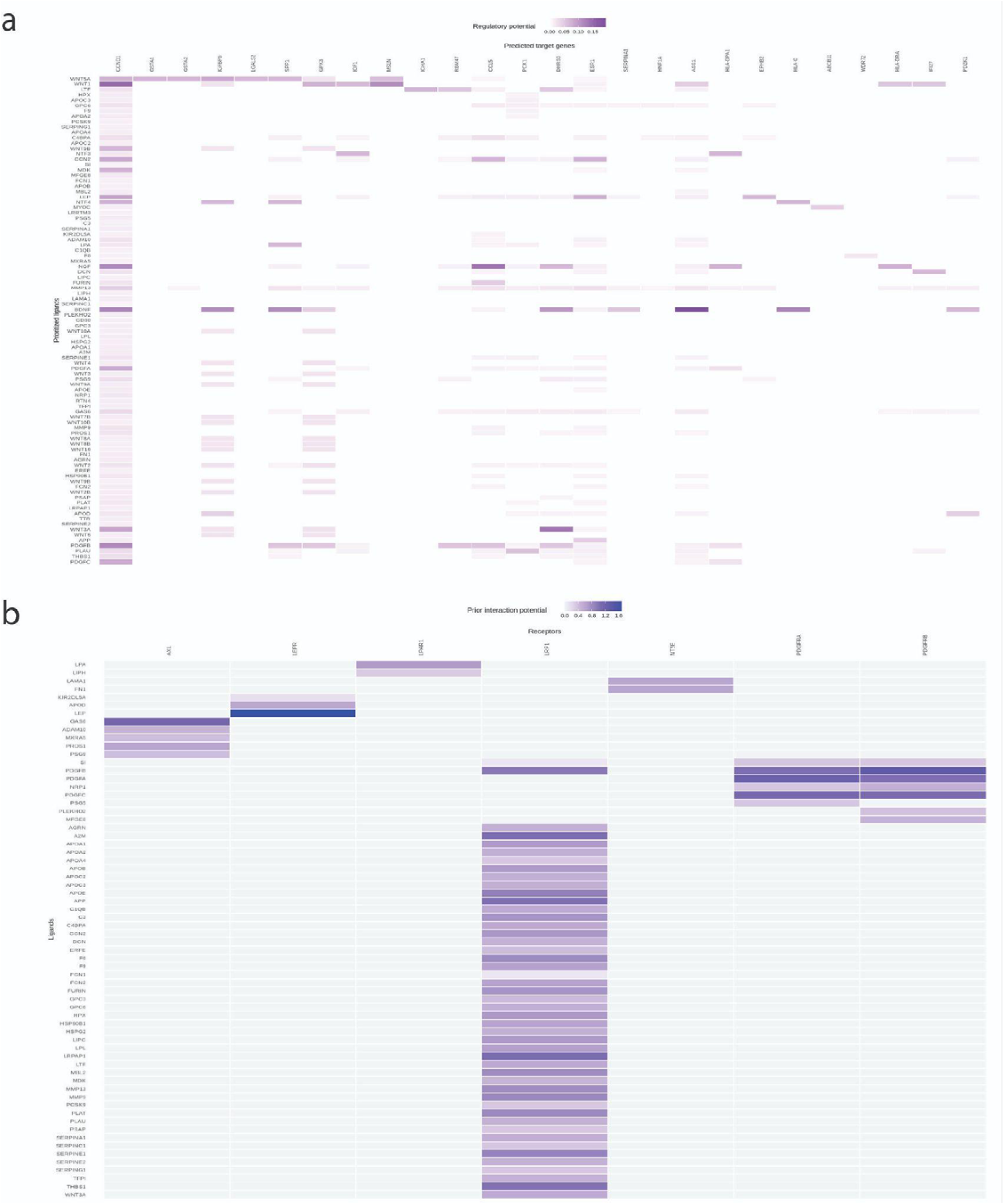
Ligand-receptor interactions and regulatory potential in the Norn cell niche. a) Ligand-target regulatory potential. Heatmap representing the predicted influence of prioritized ligands (y-axis) on the expression of target genes (x-axis) in Norn cells. High scores (dark purple) indicate strong evidence for the regulation of Norn-specific markers and survival genes—such as *CCND1, IGFBP6,* and *GPX3*—by signaling families including *WNT* and *NGF/BDNF*. b) Prior interaction potential. Heatmap of prioritized ligands and their cognate receptors identifying the most likely physical signaling axes in the Norn cell niche. High potential for PDGFB/C-PDGFRB and PDGFA-PDGFRA interactions confirms the identity of Norn cells within the PDGFR-responsive stromal lineage, while the prominent scores for the LEP-LEPR (Leptin) and GAS6-AXL axes suggest a role for systemic metabolic sensing and anti-apoptotic signaling in maintaining Norn cell homeostasis.

**Extended Figure 15.**
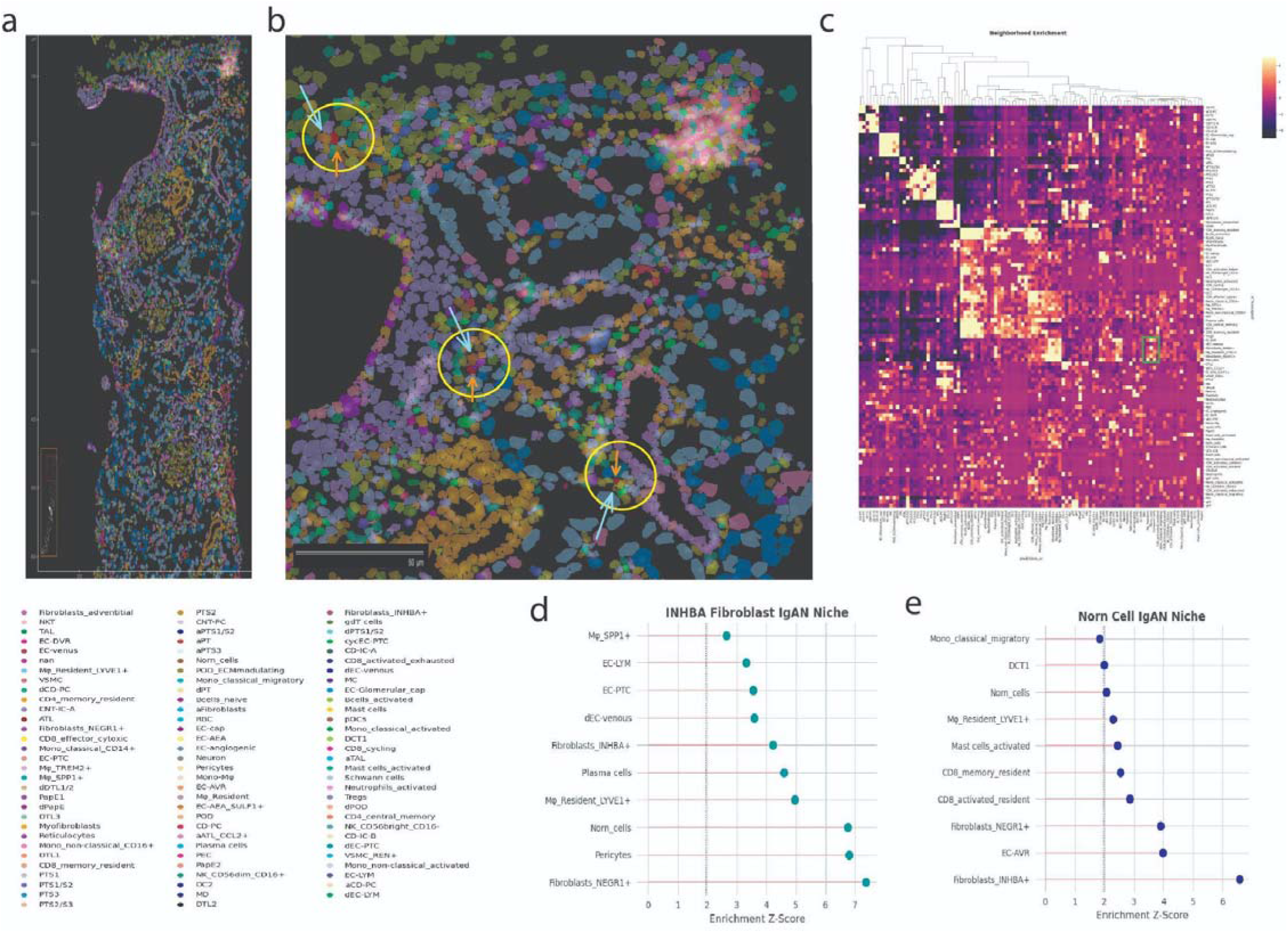
Spatial niche remodeling and Norn cell displacement in IgA Nephropathy. a) Spatial label transfer and cell type mapping. Xenium in situ hybridization (left) and high magnification inset (middle) showing the renal cortex of an IgA nephropathy (IgAN) patient. Cell types were identified via optimal transport-based label transfer from the HKCA. Yellow circles and arrows highlight the spatial co-localization of Norn cells (red) with activated INHBA^+^ (blue) populations within the fibrotic interstitium. b) Neighborhood enrichment analysis of the IgAN interstitium. Heatmap (right) displaying Z-scores for spatial proximity between all identified cell types. Hierarchical clustering reveals distinct "niche modules," with a prominent cluster (green box) identifying a recurrent association between Norn cells, INHBA^+^ fibroblasts, and NEGR1^+^ immune attracting population. c-d) Segmented niche enrichment profiles for INHBA^+^ Fibroblasts and Norn cells. Lollipop plots showing the top enriched neighbors (Z-score) for INHBA^+^ Fibroblasts (left) and Norn cells (right). The INHBA+ Niche shows high-confidence spatial association with NEGR1^+^ fibroblasts, Pericytes, and Norn cells, suggesting a consolidated activated-stromal unit. In the IgAN environment, there is spatial rearrangement correlating with areas of interstitial expansion and reflects the recruitment of Norn cells into pro-fibrotic signaling hubs.

**Extended Figure 16.**
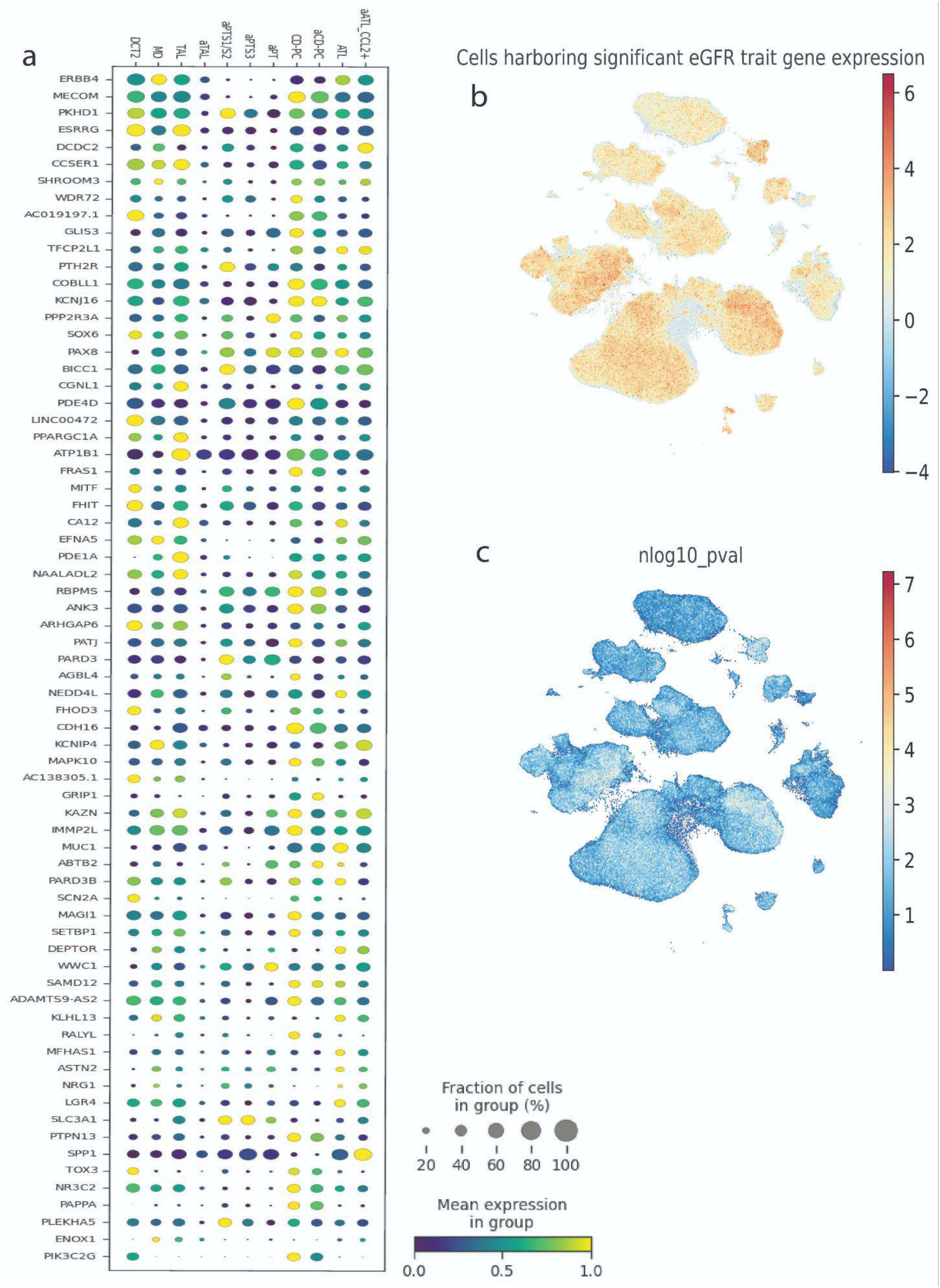
Single-cell mapping and expression profiling of eGFR trait-associated genetic risk. a) Dot plot displaying the expression of the top eGFR/creatinine trait-associated genes across distinct healthy and altered nephron loop epithelial populations. The size of each dot represents the fraction of cells within the specific cluster expressing the gene, while the color gradient indicates the mean normalized expression level within the expressing cells. b) Uniform Manifold Approximation and Projection (UMAP) of the single-cell kidney atlas, with cells colored by their normalized single-cell disease relevance score (scDRS) for the eGFR trait. Warmer colors (red) indicate cells with a higher aggregate expression of polygenic risk genes. c) UMAP embedding colored by the corresponding log10p-value of the scDRS scores, highlighting the statistical significance of trait enrichment at single-cell resolution.

**Extended Figure 17.**
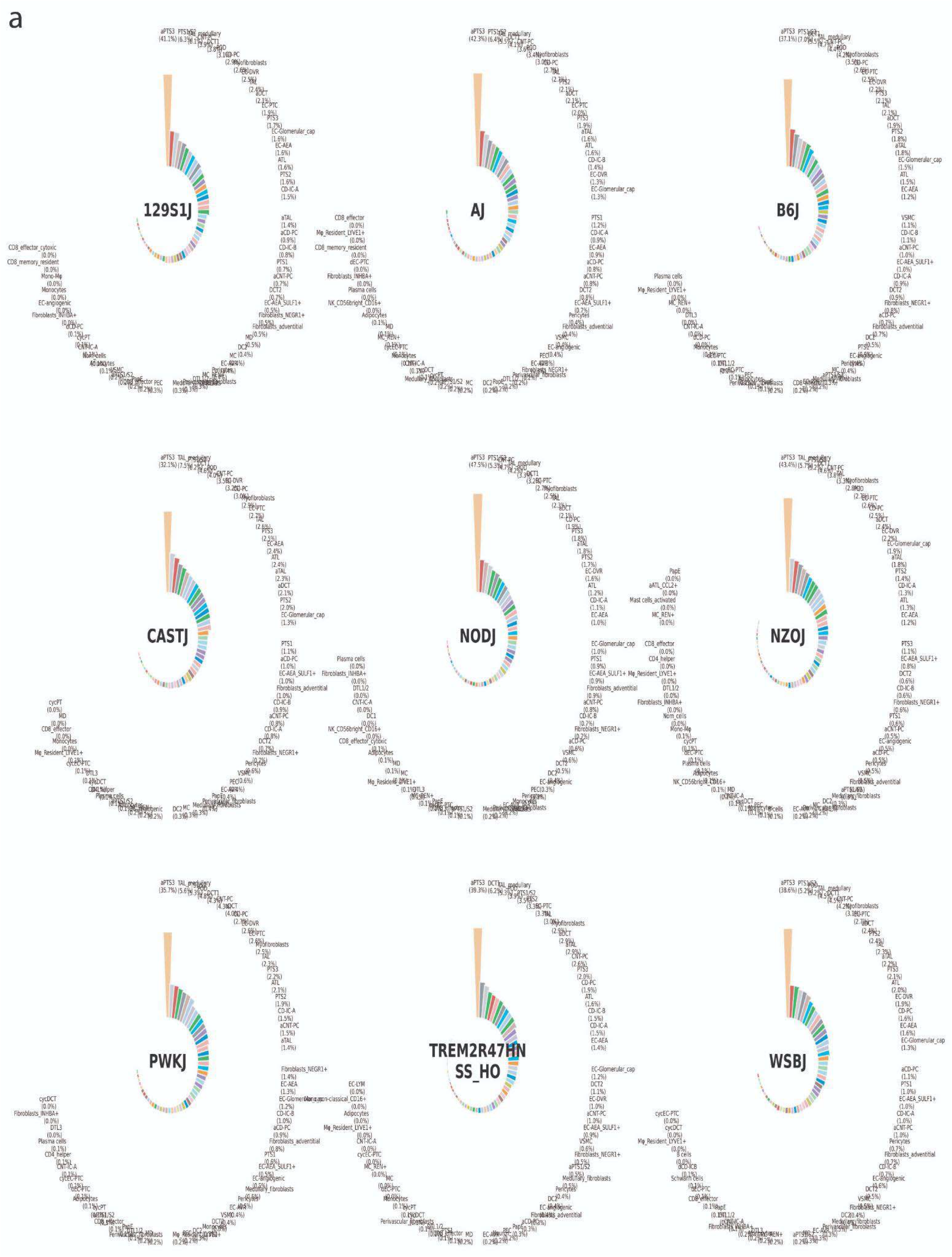
Proportions of predicted cell types amongst murine strains. The demultiplexed post-processed kidney subpools were downloaded from the IGVF website. After initial QC, 40,000 nuclei underwent HKCA model prediction. The cross-strain comparison showed aPTS3 to be the most abundant cell type across strains suggesting maintenance of an adaptive program in murine strains. We identified a much better representation of the medullary TAL in accordance with the increased availability for medullary samples.

